# GEARBOCS: An Adeno Associated Virus Tool for *In Vivo* Gene Editing in Astrocytes

**DOI:** 10.1101/2023.01.17.524433

**Authors:** Dhanesh Sivadasan Bindu, Justin T. Savage, Nicholas Brose, Luke Bradley, Kylie Dimond, Christabel Xin Tan, Cagla Eroglu

## Abstract

CRISPR/Cas9-based genome engineering enables rapid and precise gene manipulations in the CNS. Here, we developed a non-invasive astrocyte-specific method utilizing a single AAV vector, which we named GEARBOCS (Gene Editing in AstRocytes Based On CRISPR/Cas9 System). We verified GEARBOCS’ specificity to mouse cortical astrocytes and demonstrated its utility for three types of gene manipulations: knockout (KO); tagging (TagIn); and reporter knock-in (GeneTrap) strategies. Next, we deployed GEARBOCS in two test cases. First, we determined that astrocytes are a necessary source of the synaptogenic factor Sparcl1 for thalamocortical synapse maintenance in the mouse primary visual cortex. Second, we determined that cortical astrocytes express the synaptic vesicle associated Vamp2 protein and found that it is required for maintaining excitatory and inhibitory synapse numbers in the visual cortex. These results show that the GEARBOCS strategy provides a fast and efficient means to study astrocyte biology *in vivo*.

**Motivation:** Astrocytes are indispensable for brain development, function, and health. However, molecular tools to study astrocyte biology and function *in vivo* have been largely limited to genetically modified mice. Here, we developed a CRISPR/Cas9-based gene editing strategy within a single AAV vector that enables efficient genome manipulations in astrocytes. We designed and optimized this easy-to-use viral tool to understand gene expression, protein localization and function in astrocytes *in vivo*.

## Introduction

CRISPR-Cas9 (i.e., <u>C</u>lustered regularly interspaced <u>s</u>hort <u>p</u>alindromic repeats-associated endonuclease 9) mediated gene-editing is widely used to engineer the mammalian genome ^1–4^. Cas9 cleaves the genome at specific guide RNA (gRNA) target sites generating double-strand breaks. These breaks are repaired by two cell-intrinsic repair mechanisms: non-homologous end-joining (NHEJ) or homology-directed repair (HDR)^5,6^. Cas9-mediated cleavage events continue until repair induces insertions or deletions in the genome (i.e., indels) precluding gRNA recognition and further cleavage by Cas9. Indels within coding sequences often result in frameshifts and premature stop codons generating gene knockouts. HDR and NHEJ following CRISPR-Cas9-mediated genome cleavage can also be used to introduce engineered sequences into the genome, even in postmitotic cells such as neurons^7,8^. These knock-in strategies utilize in-frame donor sequences to insert reporters or tags within the open reading frames of target genes.

Numerous methods, such as SLENDR^9^, vSLENDR^10^, HITI^11^, HiUGE^12^, and ORANGE^13^ were developed for gene knockout and knock-in applications in neurons, each using different DNA delivery methods or molecular strategies for donor insertion. For example, HDR-based SLENDR and vSLENDR use *in utero* electroporation or Adeno-associated virus (AAV)-based methods to deliver gene-editing machinery into the mitotic radial glial stem cells or postmitotic neurons^9,10^. However, the efficiency of HDR-mediated donor insertion in postmitotic cells is limited^9^. The homology-independent targeted integration (HITI) method offers more effective genome editing and endogenous tagging of proteins of interest in postmitotic cells such as neurons^11^. Homology-independent Universal Genome Engineering (HiUGE) utilizes universal guide/donor pairs to enable rapid alterations of endogenous proteins without the need for designing target-specific donors^12^. Even though these gene-editing tools have versatile *in vitro* and *in vivo* applications, they depend on multiple vectors harboring the essential components required for genome engineering. Moreover, these strategies are developed primarily for neuronal gene manipulation.

In the CNS, non-neuronal glial cells, called astrocytes, play critical roles in neuronal connectivity and health. Astrocytes are characterized by their complex, ramified structures which are arborized into several thin filamentous branches^14,15^. Astrocytes are indispensable for the functioning of the brain. They provide trophic support to neurons, maintain the blood-brain barrier, regulate ion/water homeostasis as well as control synaptic transmission^16–19^. These essential roles of astrocytes in synapse formation, maturation, elimination, and maintenance depend on a complex bidirectional interaction with neurons^20,21^ mediated by secreted and adhesion molecules^22,23^.

Despite the undeniable importance of astrocytes in brain development and function, our molecular understanding of this glial cell type is still in its infancy. To address this knowledge gap, many recent studies revealed astrocyte-specific gene expression profiles and proteomes in the mammalian brain both in normal and disease conditions^24–32^. These -omics approaches provide a rich resource for future discoveries; however, efficient tools for rapid *in vivo* genome editing of mouse astrocytes are needed to speed up discovery.

Here, we developed an astrocyte-specific CRISPR-based gene-editing tool, GEARBOCS (<u>G</u>ene <u>E</u>diting in <u>A</u>st<u>R</u>ocytes <u>B</u>ased <u>O</u>n <u>C</u>RISPR/Cas9 <u>S</u>ystem) to address this need. GEARBOCS is a single AAV vector to be used in conjunction with transgenic mice expressing Cas9 in a Cre-dependent manner. GEARBOCS enables three genome engineering strategies: 1) Gene Knockout (KO), 2) knocking in a tag for labeling of endogenous proteins (TagIn) and, 3) simultaneous KO and reporter knock-in (GeneTrap). Importantly, GEARBOCS harbors all essential components for genome editing in a single AAV backbone. We tested the efficiency and specificity of GEARBOCS for *in vivo* gene editing and applied it to two experimental questions: 1) to determine if astrocytes are a required source of the synaptogenic factor Sparcl1 to maintain thalamocortical synapses after the initial synaptogenic period is completed in the mouse visual cortex and 2) to determine whether astrocytes express the synaptic vesicle associated protein Vamp2 and whether astrocytic Vamp2 is required for maintenance of synapse numbers.

Sparc-like-1 (Sparcl1, also known as Hevin) is a member of the secreted protein acidic, rich in cysteine (SPARC) family of proteins. Previous studies identified Sparcl1 as an astrocyte-secreted protein which promotes synaptogenesis^33^. Sparcl1 bridges neuronal presynaptic neurexin-1alpha and postsynaptic neuroligin-1B and orchestrates the formation of thalamocortical synapses in the developing visual cortex^34,35^. Previous studies utilized a global Sparcl1 knockout mouse to show that astrocyte-specific overexpression of Sparcl1 was sufficient to rescue the decreases in thalamocortical synapse number and ocular dominance plasticity observed in Sparcl1 KO animals^34^. However, the impact of astrocyte-specific loss of Sparcl1 on thalamocortical synapses has not been tested.

Astrocytes are known to secrete many neuroactive proteins and small molecules; however, the secretory mechanisms in these cells are not well known. Vamp2 (also known as synaptobrevin II) is a component of the soluble NSF attachment protein receptor (SNARE) complex, and its functions at neuronal synapses for regulation of neurotransmitter release are well-characterized^30,36,37^. However, the expression and functional relevance of Vamp2 and the role of the SNARE complex in astrocytes has been controversial^38,39^. Therefore, here we used GEARBOCS to determine the expression of Vamp2 in astrocytes *in vivo* and tested its roles in controlling synapse numbers in the mouse cortex.

## Results

### GEARBOCS is designed as a single AAV tool for *in vivo* genome editing in astrocytes

To facilitate rapid, astrocyte-specific, *in vivo* genome editing, we designed a CRISPR tool within a single AAV vector, which we named GEARBOCS. GEARBOCS vector is designed to be used in combination with an AAV capsid such as PHP.eB^40^ which is delivered into the CNS of mice via retro-orbital injection. For the GEARBOCS strategy to work, transgenic mice that express spCas9 in a Cre recombinase-dependent manner are required **(Figure 1A).** For all the experiments described here, we used GEARBOCS AAVs produced with AAV-PHP.eB capsids and retro-orbitally injected them into the loxP-STOP-loxP spCas9 (LSL-Cas9) or loxP-STOP-loxP spCas9-EGFP (LSL-Cas9-EGFP) mice at postnatal day (P) 21 **(Figure 1A)**. 3 weeks later, at P42, brains were harvested and analyzed. For consistency, we focused our analyses on the primary visual cortex (V1).

**Figure 1:**
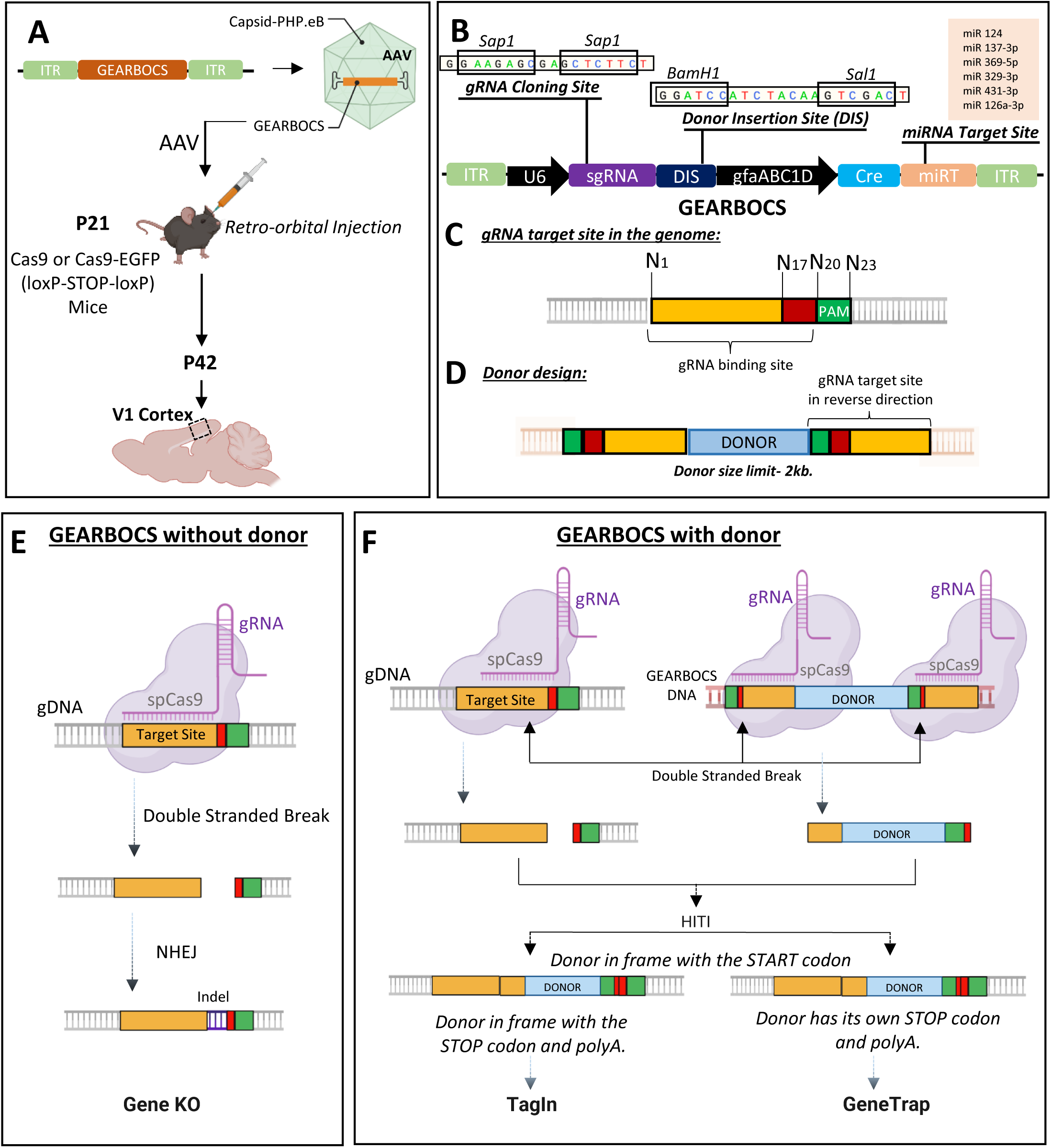
Development of GEARBOCS and its Applications: **A)** Experimental scheme for GEARBOCS-mediated *in vivo* genome editing in cortical astrocytes. GEARBOCS AAVs are produced with AAV-PHP.eB capsid and retro-orbitally injected into the loxP-STOP-loxP Cas9 or Cas9-EGFP mice at P21. Brains are prepared for immunohistochemistry and subsequent confocal fluorescent imaging at P42. **B)** Schematic of GEARBOCS vector showing its four essential elements for CRISPR/Cas9-based genome editing: 1) A human U6 promoter that can drive the expression of a unique guide RNA (gRNA) which is cloned into the gRNA cloning site (*Sap1* restriction enzyme); 2) Donor Insertion Site (DIS) wherein the donor DNA (e.g. encoding epitope tags, EGFP, mCherry etc.) can be cloned between the BamH1 and Sal1 restriction sites, 3) Cre expression cassette driven by astrocyte-specific gfaABC1D promoter with microRNA targeting cassette; 4) AAV2 Inverted terminal repeats (ITR) for AAV packaging and expression. **C).** Guide RNA target site present in the genome is recognized by the unique gRNA/Cas9 complex through its recognition sequence and PAM sequence (green box) followed by the double stranded break at cut site (between red and yellow box). **D).** Donor DNA designed to clone into the GEARBOCS has the donor sequence (Light blue Box) flanked by the guide RNA target sites at both ends. GEARBOCS can accommodate up to 2kb size donors for AAV mediated cargo delivery and *in vivo* genome editing. **E-F** Schematic of GEARBOCS mediated genome editing mechanism in astrocytes. In GEARBOCS without donor model, AAV mediated delivery of GEARBOCS into the mouse cortex in loxP-STOP-loxP Cas9 mice leads to the Cre-mediated expression of spCas9 in astrocytes through the gfaABC1D promoter in GEARBOCS. **E**) In the absence of a donor sequence, U6 driven gRNA, and spCas9 cause double strand breaks in the gene of interest, which is followed by the imprecise non-homologous end joining (NHEJ) repair process, leading to indels and subsequent gene knockout (KO). **F)** In GEARBOCS with donor, guided by the gRNA, spCas9 makes double stranded breaks both within the gene of interest and the GEARBOCS vector around the donor sites. The excised donor fragment from the GEARBOCS can integrate into the genome by homology-independent targeted integration (HITI). The donors are designed to be in frame with the gene of interest. If the endogenous tagging of a protein of interest in astrocytes (TagIn) is needed, the tag is knocked in frame with both the START and STOP codons of the gene of interest. If the donor has its own STOP codon and polyA tail, then this will lead to GeneTrap.

The GEARBOCS vector (5743bp) harbors four essential components for *in vivo* genome editing:

1. A U6 expression cassette, including a human U6 promoter, a gRNA cloning site, and a 76bp gRNA scaffold^41^. The U6 promoter drives the expression of a small guide RNA (gRNA), which is cloned into the *Sap1* restriction site within the gRNA cloning site **(Figure 1B)**. The gRNA is designed to match a 20bp genomic target site, which is next to a Protospacer Adjacent Motif (PAM) **(Figure 1C)**.
2. A Donor Insertion Site (DIS) wherein the donor DNA can be cloned between the BamH1 and Sal1 restriction sites **(Figure 1B)**. The donor DNA is omitted for generating gene KOs. Alternatively, donor sequences, ranging from a small epitope tag, like HA or V5, or a larger reporter tag, such as EGFP or mCherry (up to 2kb), can be cloned into the DIS. The donor sequence is flanked with the guide RNA target site sequences on both ends **(Figure 1D)**.
3. A synthetic human glial fibrillary acidic protein (GFAP) promoter^42^, GfaABC1D, to drive the expression of Cre in astrocytes. Cre removes the STOP codon and turns on Cas9 or Cas9-EGFP expression in astrocytes. While the GfaABC1D promoter is highly active in astrocytes, only low levels of cre-recombinase expression are necessary for recombination. As a result, an early version of GEARBOCS (GEARBOCS-v0) led to recombination in a small number of neurons^43^ (**Figure S1**). To overcome this, we incorporated a miRNA targeting cassette (miRT) to the 3 prime of the Cre-recombinase sequence. This miRNA targeting cassette triggers transcript degradation in neurons and endothelial cells, increasing astrocyte specificity^44^.
4. Two inverted terminal repeats (ITRs) for AAV packaging. The ITR sequences in GEARBOCS are based on the AAV2 serotype^45^.

The GEARBOCS approach uses the intrinsic DNA repair mechanism non-homologous end joining (NHEJ) for homology-independent targeted integration (HITI). Thus, when combined with HITI of donor sequences^11^, GEARBOCS permits gene knock-in both in mitotic and postmitotic cells. The GEARBOCS tool can be used to manipulate the astrocyte genome in three different ways. GEARBOCS, without a donor **(Figure 1E)**, generates knock out (KO) of the gene of interest in astrocytes. The gRNA specific to the gene of interest guides the spCas9 in astrocytes to the genomic locus to create a double stranded break upstream of its PAM sequence (NGG for spCas9). This double stranded break is repaired by NHEJ, until the repair causes indels which results in a frame shift and gene KO. To generate KOs, we choose unique gRNAs targeting genomic regions within the first few exons of the gene of interest. gRNAs are chosen based on their low off-target and high on-target scores predicted by the Broad Institute CRISPick tool (available at https://portals.broadinstitute.org/gppx/crispick/public)^46,47^.

GEARBOCS can also be used for knocking in a donor sequence in frame with the gene of interest **(Figure 1F)**. To do so, a donor sequence, flanked with the reverse-oriented genomic gRNA target sites on both ends, is cloned into the DIS of the GEARBOCS vector **(Figure 1B and D)**. The gRNAs guide spCas9s to the genomic loci and the donor sequence within the viral genome to generate double-stranded breaks in both sequences. These cleavage events result in two blunt ends in the genome and two at the donor fragment **(Figure 1F)**. The donor DNA is then integrated into the genomic target site via HITI^11^.

GEARBOCS with donor can be used for endogenous tagging of proteins of interest (TagIn) in astrocytes. To do so, the donor is designed to insert a protein coding sequence in frame with the endogenous coding sequence so that the endogenous start and stop codons and the polyA tails are kept intact **(Figure 1F)**.

When the donor sequence is designed to insert in frame following the endogenous START site but has its own STOP codon and a synthetic polyA tail, this will lead to a gene trap: The donor sequence will be expressed but the target gene will be knocked out after the editing site **(Figure 1F)**. This GeneTrap approach is particularly useful to label and visualize astrocytes in which a gene of interest is knocked out. Reverse orientation of gRNA target sequences flanking the donor (**Figure 1D)** is critical for avoiding further cleavage, thereby facilitating the insertion of the donor in the correct orientation^7^. The GEARBOCS vector design enables us to interchange the donors quickly with a single cloning step. Hence, it provides an “all-in-one” simple CRISPR tool to target astrocytes *in vivo*.

### GEARBOCS enables efficient astrocyte-specific genome editing

GEARBOCS uses the astrocyte-specific minimal promoter, gfaABC1D^42^ to drive the Cre expression and turn on the production of Cas9 or Cas9-EGFP in astrocytes of the commercially available LSL-Cas9 or LSL-Cas9-EGFP mice^48,49^. This approach overcomes the difficulty of expressing large-sized Cas9 orthologs through the AAV system.

Previous studies have raised the caveat that the human GfaABC1D promoter may permit Cre-mediated genome recombination in some neuronal cells, when AAVs are directly injected at high titers into the mouse brain^50^. Thus, we first tested the astrocyte-specificity of GEARBOCS in the mouse cortex for its *in vivo* applications. To do so, we used the AAV capsid PHP.eB to package the GEARBOCS vector. This capsid efficiently transduces CNS cells^40^ when retro-orbitally injected, eliminating the need for invasive surgery and direct injection into the CNS. We found that retro-orbitally injected AAV-GEARBOCS indeed transduces the mouse brain with high efficiency which is evidenced by the abundance of Cas9-EGFP positive cells across many brain regions **(Figure 2A-B)** including V1 cortex **(Figure 2C)**

**Figure 2:**
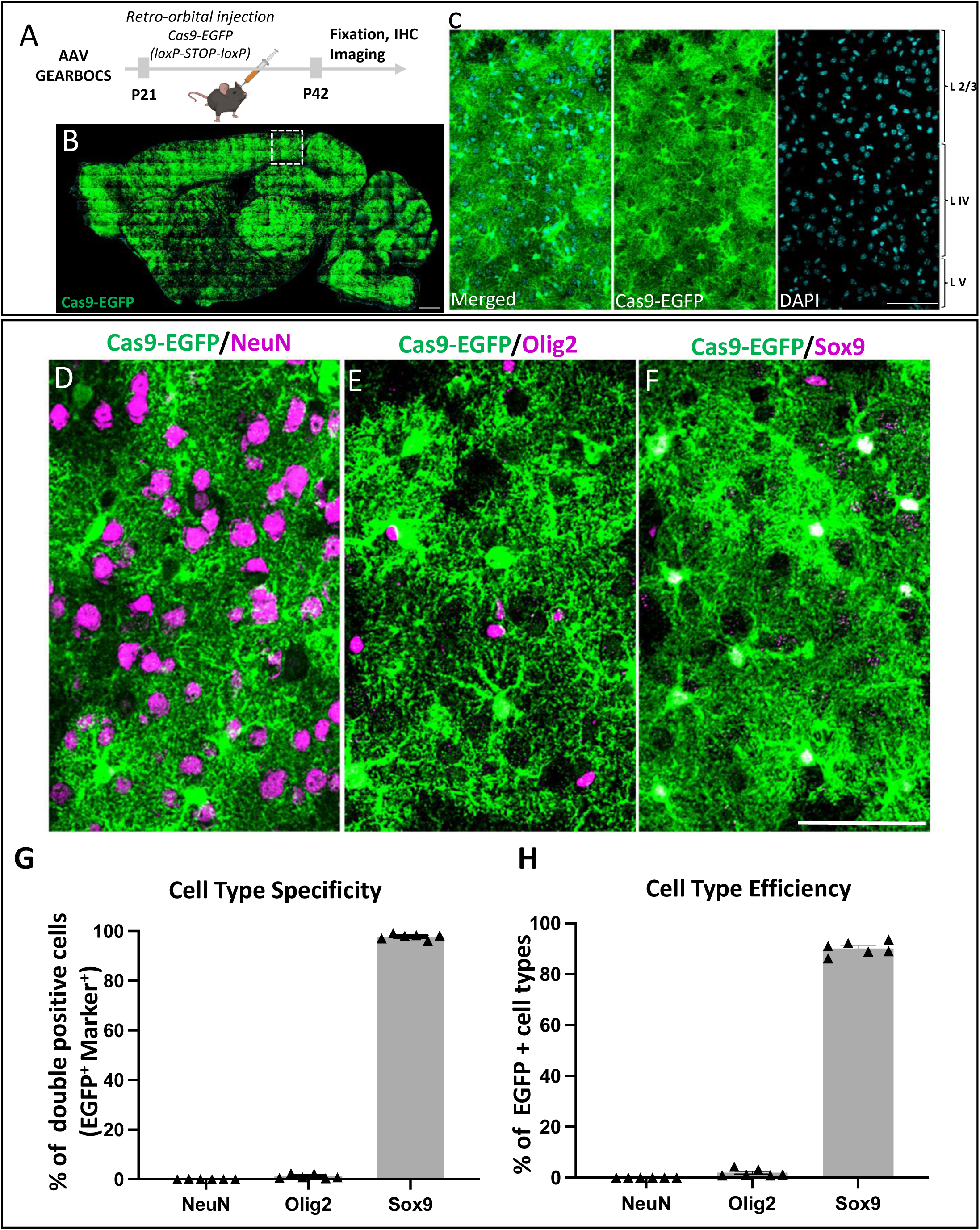
GEARBOCS efficiently targets mouse cortical astrocytes: **A).** Schematic of the experimental scheme. GEARBOCS AAV is generated with AAV-PHP.eB capsid and retro-orbitally injected into the loxP-STOP-loxP Cas9-EGFP mice at P21. Three weeks post viral injection (P42), brains are collected for analysis. **B)** Confocal immunofluorescent image of a sagittal section from AAV-GEARBOCS-injected loxP-STOP-loxP Cas9-EGFP mouse brain showing the prevalence of Cas9-EGFP expression (green). Scale bar=50μm. **C).** Confocal immunofluorescent image from V1 cortex (indicated as dotted box in **B**) showing the Cas9-EGFP positive cells in different cortical layers. Scale bar=30μm. **D-F)** Confocal immunofluorescent images showing EGFP positive cells (green) co-stained with different cell type-specific markers (magenta) such as **(D)** NeuN (neurons), **(E)** Olig2 (oligodendrocytes), and **(F)** Sox9 (astrocytes). Scale bar=20μm. **G)** Cell type-specificity of GEARBOCS-Cre expression quantified by as the percentage of each cell types co-localized with EGFP positive cells in the V1 cortex**. H)** Cell type efficiency of GEARBOCS-Cre expression quantified by taking the percentage of EGFP positive cells co-localized with cell type marker in the V1 cortex. (G-H: n=6 animals, Error bars show SEM).

To investigate the astrocyte specificity of Cre-mediated Cas9 expression, we performed the co-immunostaining of Cas9-EGFP with different CNS cell type-specific markers. We used NeuN to label neurons^51^, Olig2 for oligodendrocyte lineage cells^52^, and Sox9 that labels the majority of the astrocytes^53^ **(Figure 2D-F)**. The quantification of marker colocalization with the EGFP/Cas9-expressing (i.e., EGFP/Cas9+) cells in the cortex revealed that they were overwhelmingly Sox9+ (97.82±0.44%) compared to NeuN+ (0.035±0.035%) or Olig2+ cells (1.23±0.37%) **(Figure 2G)**. Of all the Sox9+ astrocytes, 90.15±1.07% were EGFP/Cas9+, whereas only 0.0065±0.0065% and 2.01±0.59% of NeuN+ or Olig2+ cells were EGFP/Cas9+, respectively (**Figure 2H)**. These results show that GEARBOCS targets Cas9 expression to cortical mouse astrocytes with high specificity and efficiency.

### GEARBOCS-mediated gene editing of Sparcl1 in mouse cortical astrocytes

To evaluate the effectiveness of GEARBOCS as a molecular CRISPR tool for manipulation of astrocyte genome in multiple ways, we targeted the gene Sparcl1, which encodes for a secreted synaptogenic protein, also known as Hevin^34,35,54^. Previous studies on Hevin/Sparcl1’s role in synapse formation specifically focused on early postnatal development and utilized global Sparcl1-KO mice^34,35^. Here we used GEARBOCS to target Sparcl1 specifically in astrocytes after the completion of the synaptogenic period (after P21) to determine if its astrocytic expression is required for maintenance of thalamocortical synapses. To do so, we first applied the GEARBOCS KO strategy **(Figure 1C)** by selecting a unique gRNA targeting the first exon of Sparcl1, located 40bp after the start codon (**Figure 3A**). The selected gRNA1 was cloned into GEARBOCS to generate the AAV-GEARBOCS-Sparcl1-KO and retro-orbitally injected into P21 Cas9-EGFP (loxP-STOP-loxP) mice. We used AAV-GEARBOCS without a gRNA as our control. After tissue collection at P42, immunostaining showed that the Sparcl1 expression is ablated in EGFP/Cas9+ cells which are transduced by AAV-GEARBOCS-Sparcl1-KO when compared to the astrocytes transduced by the control virus **(Figure 3B-G)**. Of the Cas9-EGFP positive astrocytes in the AAV-GEARBOCS-Sparcl1-KO condition, 95.58±0.31% were lacking Sparcl1 staining **(Figure 3H-K)**. This result shows that GEARBOCS can be used to knockout gene(s) of interest in astrocytes with high efficiency.

**Figure 3:**
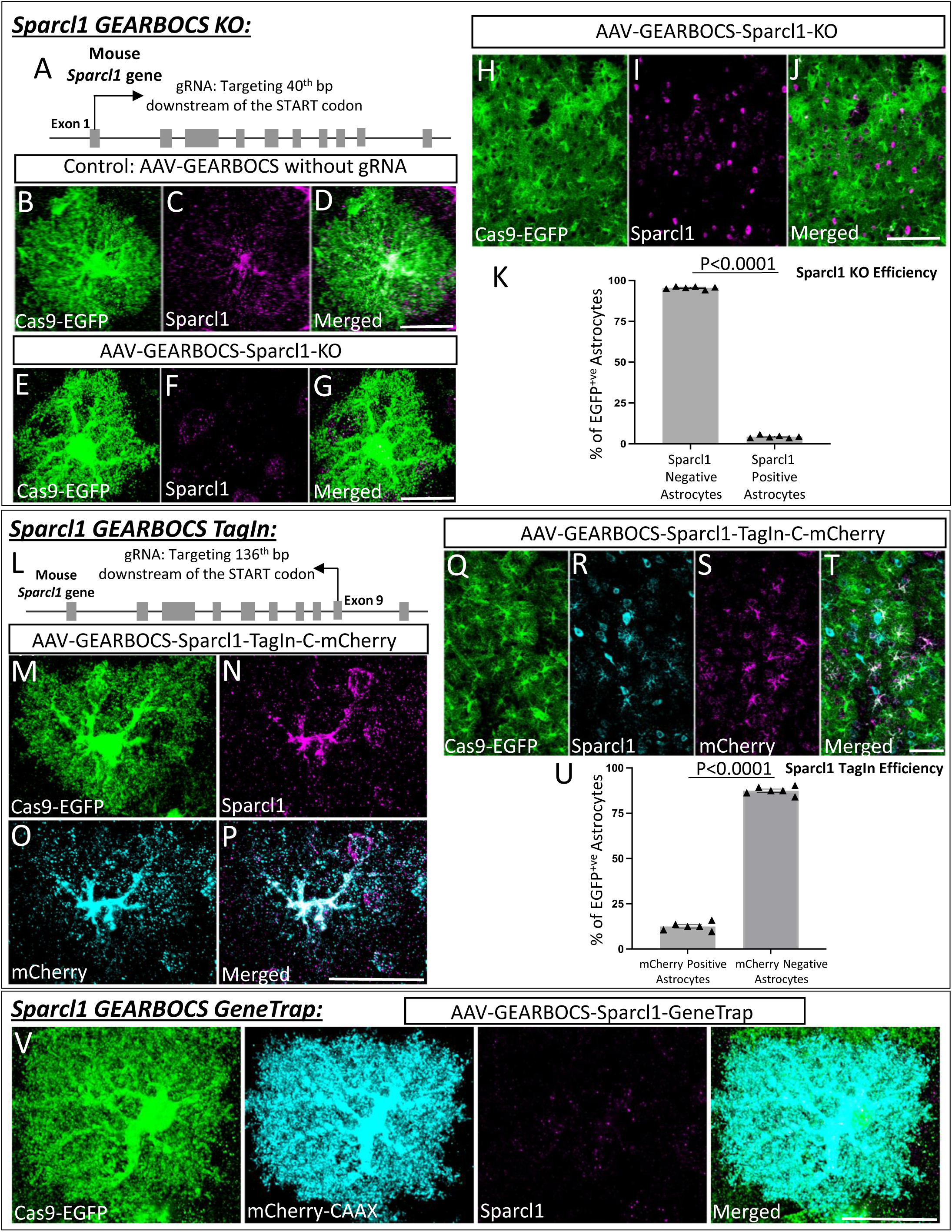
GEARBOCS-mediated editing of *Sparlc1* gene in astrocytes: **A).** Schematic showing the first gRNA target site in *Sparcl1* locus used in this study. **B-D).** Confocal immunofluorescence microscopy images showing a Cas9-EGFP positive astrocyte (green) transduced with AAV-GEARBOCS vector without the gRNA (Control). This astrocyte still has Sparcl1 staining (magenta). **E-G).** Confocal immunofluorescence microscopy images showing the loss of Sparcl1 staining (magenta) in a Cas9-EGFP (green) positive AAV-GEARBOCS-*Sparcl1*-KO transduced astrocyte. **H-J)** Confocal immunofluorescence microscopy images of a Cas9-EGFP-positive astrocytes (green) transduced with the AAV-GEARBOCS-Sparcl1-KO in a large field of view. **K)** Quantification of the percentage of Cas9-EGFP positive astrocytes that are Hevin positive or negative by immunohistochemistry. N = 6 animals, p <0.001 by student’s t-test. **L)** Schematic showing the gRNA target site in the mouse *Sparcl1* gene for AAV-GEARBOCS-Sparcl1-TagIn-C-mCherry. **M-P).** Confocal Immunofluorescence microscopy images of a Cas9-EGFP positive astrocyte (green) transduced with AAV-GEARBOCS-Sparcl1-TagIn-C-mCherry showing the co-localization of Sparcl1 (magenta) with mCherry (cyan). **Q-T).** Confocal immunofluorescence image of Cas9-EGFP positive astrocytes (green) transduced with the AAV-GEARBOCS-Sparcl1-TagIn-C-mCherry in a large field of view. **U)** Quantification of the percentage of Cas9-EGFP positive astrocytes that are mCherry positive or negative by immunohistochemistry. N = 6 animals, p <0.001 by student’s t-test. Scale bars=20μm. **V)** Confocal immunofluorescence image of Cas9-EGFP positive astrocyte transduced with the AAV-GEARBOCS-Sparcl1-GeneTrap. Scale bars=20μm.

To test the utility of GEARBOCS for TagIn applications **(Figure 1D)**, we designed a donor sequence, containing an mCherry fluorophore tag sequence, flanked with two inverted gRNA target sequences on either side. This gRNA sequence corresponded to a suitable donor insertion site in exon 9 of Sparcl1 at the C terminus of the protein. This donor sequence was then cloned into the GEARBOCS vector, which already contained the gRNA targeting the exon 9, generating AAV-GEARBOCS-Sparcl1-TagIn-C-mCherry. We validated the efficiency of the guide and potential of the donor insertion first using 3T3-Cas9 cells (Calibre Scientific, Cat# CBIO-AKR-5104), a mouse fibroblast cell line expressing spCas9. By PCR amplifying the edited genomic DNA of these cells, we were able to validate this approach (**Figure S2J**).

AAV-GEARBOCS-Sparcl1-TagIn-C-mCherry viruses were injected into the floxed-Cas9-EGFP mice and cortical sections were stained with antibodies against Sparcl1 and mCherry. As predicted, the endogenous Sparcl1 staining co-localized with mCherry in the GEARBOCS-transduced EGFP/Cas9+ astrocytes **(Figure 3M-T).** Of the Cas9-EGFP positive astrocytes, 12.49±0.90% had mCherry-tagged Sparcl1 (Figure 3U).

We were also able to label Sparcl1’s N-terminus with either mCherry or HA TagIn viruses and observed colocalization between Sparcl1 antibody staining and these N-terminal tags **(Figure S2)**.

Finally, we tested the GEARBOCS tool’s utility to achieve a GeneTrap-i.e., the simultaneous KO of an astrocytic gene of interest and insertion of a reporter gene into its locus **(Figure 1F)**. To do so, we used the gRNA targeting the *Sparcl1* gene in exon 1 **(Figure 3A)**. We designed and generated the mCherry-CAAX reporter donor with its own STOP codon and polyA tail. This GeneTrap strategy caused loss of endogenous Sparcl1 expression as expected, while the mCherry-CAAX was produced in these cells facilitating their visualization **(Figure 3V-Y)**. Collectively, these results demonstrated that GEARBOCS is an efficient tool for gene KO, endogenous tagging, and gene trap purposes, greatly facilitating the study of astrocyte cell biology in the mouse CNS. Hence, this “all-in-one” AAV CRISPR tool could help address the limitations and challenges in understanding the development and functions of astrocytes *in vivo*.

### Astrocytic Sparcl1 is required for thalamocortical synapse maintenance in the mouse visual cortex

Previous studies have shown that Sparcl1 is required for proper synapse formation in the mouse visual cortex. Moreover, specifically overexpressing Sparcl1 in astrocytes rescues synapse number deficits in Sparcl1 KO mice^34^. While these data show that astrocytic Sparcl1 is sufficient for synapse formation, it remains an open question if astrocytic Sparcl1 is necessary for thalamocortical synapse formation and maintenance. This is particularly important because a small subset of cortical neurons also express Sparcl1/Hevin protein^35^. Moreover, Sparcl1 mRNA is also present in endothelial cells^30^ and overexpressing Hevin specifically in the endothelial cells of the Hevin KO mice was able to rescue synapse phenotypes in the Sparcl1-KO mice^55^.

To test this, we used the GEARBOCS Sparcl1 KO system from Figure 3A to knockout Sparcl1 specifically in astrocytes **(Figure 4B)**. In the mouse visual cortex, ages between P21 and P42 coincide with the critical period of ocular dominance plasticity, during which synapses carrying eye-specific synaptic inputs can be remodeled drastically if one of the eyes loses vision^56^. Interestingly, we found that knocking out of Sparcl1 specifically in the astrocytes by the injection of Sparcl1 KO GEARBOCS at P21 and brain collection at P42 showed a 48% decrease in VGlut2/PSD95 synapse numbers **(Figure 4C-E).** These results show that in addition to its role during early synaptogenic period (before P21) astrocytic Sparcl1 is required for the maintenance of thalamocortical connectivity during the critical period of ocular dominance plasticity.

**Figure 4:**
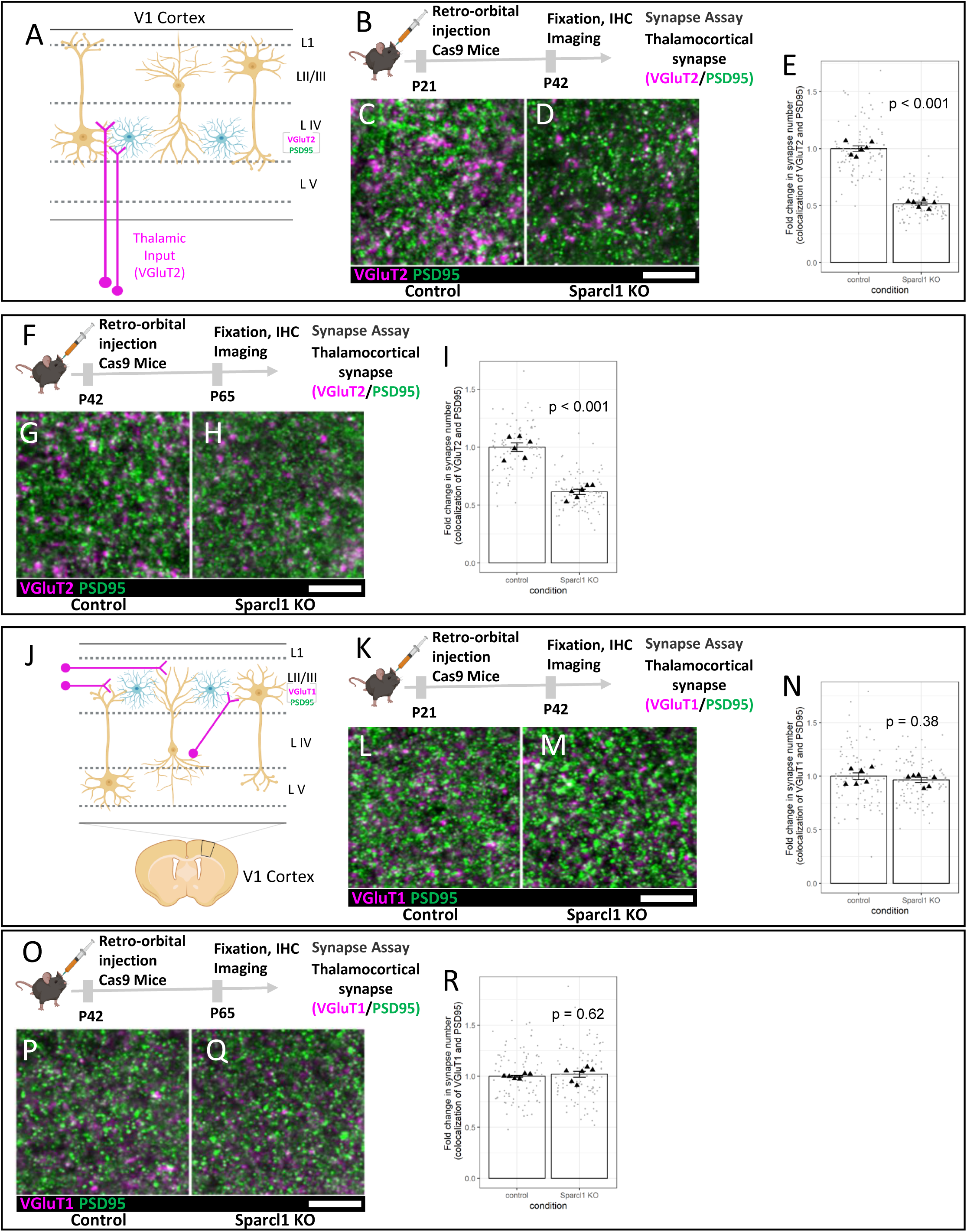
Astrocytic Sparcl1 is required for formation and maintenance of thalamocortical synapses: **A)** Schematic of thalamocortical inputs into the primary visual cortex (V1). Thalamocortical inputs (magenta) are labeled with VGlut2. **B)** Schematic of AAV-GEARBOCS-Sparcl1-KO experimental scheme. AAVs were retro-orbitally injected into the loxP-STOP-loxP Cas9 mice at P21 and the brains were collected at P42. **C-D)** Confocal immunofluorescence images of thalamocortical excitatory synapses marked as close apposition of VGluT2 (magenta) and PSD95 (green)**. E)** Quantification of synaptic density of thalamocortical co-localized puncta. **F)** Schematic of AAV-GEARBOCS-Sparcl1-KO experimental scheme. AAVs were retro-orbitally injected into the loxP-STOP-loxP Cas9 mice at P42 and the brains were collected at P65. **G-H)** Confocal immunofluorescence images of thalamocortical excitatory synapses marked as close apposition of VGluT2 (magenta) and PSD95 (green)**. I)** Quantification of synaptic density of thalamocortical co-localized puncta. **J)** Schematic of intracortical synapses in the primary visual cortex (V1). Intracortical inputs (magenta) are labeled with VGlut1. **K)** Schematic of AAV-GEARBOCS-Sparcl1-KO experimental scheme. AAVs were retro-orbitally injected into the loxP-STOP-loxP Cas9 mice at P21 and the brains were collected at P42. **L-M)** Confocal immunofluorescence images of thalamocortical excitatory synapses marked as close apposition of VGluT1 (magenta) and PSD95 (green)**. N)** Quantification of synaptic density of thalamocortical co-localized puncta. **O)** Schematic of AAV-GEARBOCS-Sparcl1-KO experimental scheme. AAVs were retro-orbitally injected into the loxP-STOP-loxP Cas9 mice at P42 and the brains were collected at P65. **P-Q)** Confocal immunofluorescence images of thalamocortical excitatory synapses marked as close apposition of VGluT1 (magenta) and PSD95 (green)**. R)** Quantification of synaptic density of thalamocortical co-localized puncta. n = 6 animals per condition. p values were calculated using a linear mixed effects model to account for the multiple images taken from each animal. Error bars represent 1 standard error of the mean. Scale bars=10μm.

To test if Sparcl1 is required for synapse maintenance after the critical period has ended, we injected Sparcl1 KO GEARBOCS at P42 and assessed VGlut2/PSD95 synapse counts at P65 (**Figure 4F**). These experiments revealed a 38% reduction in VGlut2/PSD95 synapse numbers (**Figure 4G-I**). To determine if the effect of Sparcl1 loss on thalamocortical synapses was specific or due to a general impact on synapses in the cortex, we also assessed the density of VGlut1/PSD95 intracortical synapses using the same Sparcl1 KO paradigm from 4B and 4F. Loss of Sparcl1 from astrocytes did not alter VGlut1/PSD95 synapse density in either the P21 injection, P42 collection (**Figure 4J-N**) or P42 injection, P65 collection experiments (**Figure 4O-R**). These experiments, taken together with previously published studies^34,35^, show that astrocytic Sparcl1 is required for thalamocortical synapse formation and maintenance.

### Mouse cortical astrocytes express Vamp2 *in vivo*

Astrocytes secrete synapse-modulating proteins, peptides, and neuroactive small molecules^57–59^ however, the mechanisms underlying their release are unknown. Vamp2 is an integral component of the SNARE complex which mediates calcium-dependent vesicular exocytosis^36,37^. In response to an increase in cytosolic calcium levels, vesicular Vamp2 forms ternary SNARE complexes with plasma membrane proteins such as syntaxin and SNAP23 to cause membrane fusion and release of vesicular cargo **(Figure 5A).**

**Figure 5:**
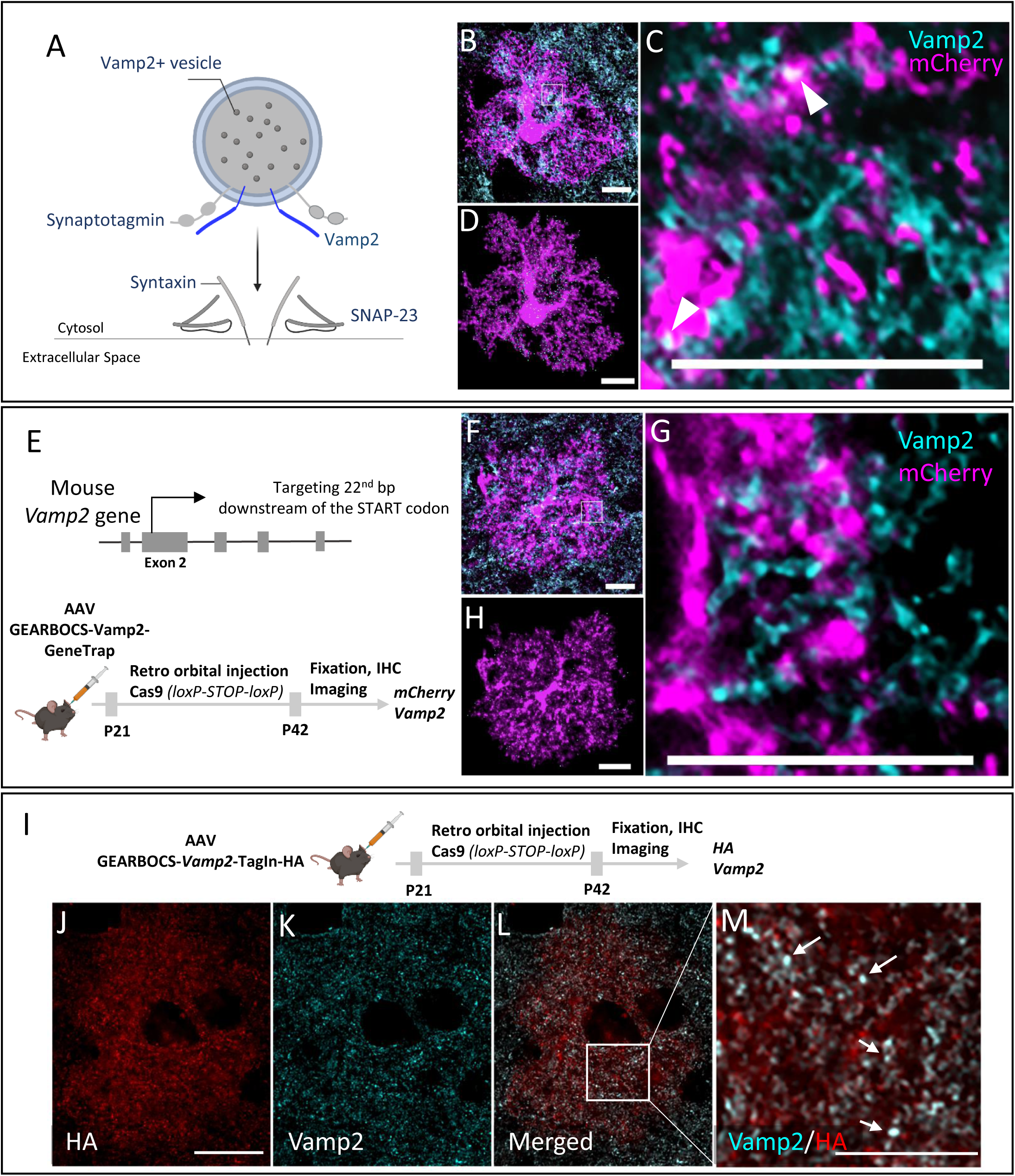
*In vivo* mouse cortical astrocytes express Vamp2: **A).** Schematic of Vamp2 (dark blue) as a member of the SNARE complex. **B)**. Confocal immunofluorescence microscopy image from an mCherry-positive astrocyte (magenta) transduced with AAV-gfaABC1D-mCherry-CAAX co-stained with a Vamp2 antibody (cyan). Scale bar=10μm. **C).** Zoom in image of a single optical section showing the presence of Vamp2 (cyan) inside the mCherry-CAAX filled astrocytes (magenta). Scale bar=10μm. **D).** IMARIS reconstructed image showing the spatial distribution of Vamp2 (cyan) within the mCherry-CAAX filled astrocyte domain (magenta); Scale bar-10μm. **E).** Schematic of mouse *Vamp2* gene showing the gRNA target site and the experimental scheme showing the GEARBOCS-*Vamp2*-GeneTrap. AAV-GEARBOCS-*Vamp2*-GeneTrap was retro-orbitally injected into the loxP-STOP-loxP Cas9 mice at P21 and immunohistochemical analyses of mCherry and Vamp2 were carried out at P42. **F).** Confocal immunofluorescence image of an mCherry positive astrocyte (magenta) from AAV-GEARBOCS-*Vamp2*-GeneTrap transduced brain showing that Vamp2 staining within the mCherry-positive astrocyte is diminished. Scale bar=10μm. **G).** Zoom in image of a single optical section showing the reduction of Vamp2 staining inside the mCherry-CAAX positive GEARBOCS-*Vamp2*-GeneTrap astrocyte (magenta). Scale bar-10μm. **H).** IMARIS reconstructed image showing the decreased expression of Vamp2 (cyan) in a GEARBOCS-*Vamp2*-GeneTrap astrocyte (magenta). Scale bar-10μm. **I).** Schematic of the experimental scheme showing the GEARBOCS-*Vamp2*-TagIn with an HA tag. AAV-GEARBOCS-*Vamp2*-TagIn-HA was retro-orbitally injected into the loxP-STOP-loxP Cas9 mice at P21 and immunohistochemical analyses of HA and Vamp2 were carried out at P42. **J-L).** Confocal immunofluorescence image of a single optical section from an astrocyte transduced with the AAV-GEARBOCS-*Vamp2*-TagIn-HA showing the co-localization Vamp2 (cyan) with HA (Red). Scale bar-20μm. **M).** Zoom in image of a single optical section showing the co-localization of Vamp2 with HA (white). Scale bar=10μm.

While the presence of calcium transients in astrocytes has been well documented^60^, the expression of Vamp2 in astrocytes and its functions are still controversial. This is primarily due to the overwhelming abundance of neuronal Vamp2 compared to astrocytic Vamp2 and the limited specificity of both chemical^39,61^ and genetic approaches to target astrocytic Vamp2^38^. Therefore, it has been difficult to discriminate the astrocytic expression of Vamp2 from its neuronal counterpart.

To capture astrocytic Vamp2 expression by conventional immunohistochemical methods and imaging, we immunostained Vamp2 in brain sections from mice in which cortical astrocytes were transduced with AAV-gfaABC1D-mCherry-CAAX. Co-localization of mCherry+ astrocytes with Vamp2 was detected; however, due to the intense Vamp2 staining in brain tissue and the resolution limit of light microscopy, it was difficult to confirm Vamp2 expression within astrocytes **(Figure 5B-D).** Using the Imaris software, we reconstructed the fluorescence signal from an mCherry-filled astrocyte to map the spatial Vamp2 expression in astrocytes **(Figure 5D).**

To verify the specificity of observed astrocytic Vamp2 staining with GEARBOCS, we used a Vamp2-targeting gRNA which was previously described^62^. Using the GeneTrap method, we knocked-in an mCherry-CAAX donor with a STOP codon and polyA tail at its second exon, 22bp after the start codon **(Figure 5E)**. This strategy allowed us to confirm the endogenous Vamp2 promoter activity driving mCherry-CAAX expression, while simultaneously knocking out Vamp2 expression. Immunohistochemical analysis of mCherry and Vamp2 in cortical astrocytes transduced with AAV-GEARBOCS-Vamp2-GeneTrap revealed greatly diminished Vamp2 staining within the mCherry+ astrocytes **(Figure 5F-H)**. These results show that Vamp2 locus is transcriptionally active in astrocytes, further suggesting that astrocytes express Vamp2.

To further corroborate this finding, we applied the GEARBOCS TagIn strategy to insert an HA epitope tag at the N-terminus of the Vamp2 protein. To do so, we used the same gRNA **(Figure 5E),** but changed our donor with an in-frame sequence for a single HA-epitope tag. We initially validated this approach by transfecting 3T3-Cas9 cells with this construct and then conducting Sanger sequencing of the PCR amplified target site. This sequencing confirmed the insertion of the proper HA or mCherry donors **(Figure S3)**. AAV-GEARBOCS-Vamp2-TagIn-HA viruses were retro-orbitally injected into the Cas9 mice at P21 **(Figure 5I)** and brains were harvested at P42. HA-tagging revealed punctate staining within transduced astrocytes, suggesting a vesicular localization **(Figure 5J).** Coimmunostaining of astrocytes with HA and Vamp2 showed that Vamp2 signal colocalizes with the HA **(Figure 5J-M)**. Altogether, these data show that cortical astrocytes express Vamp2 *in vivo.* Furthermore, these results demonstrate GEARBOCS as a powerful tool for specifically characterizing the localization and function of astrocytic proteins while leaving neuronal protein unaffected and unlabeled.

### Astrocytic Vamp2 is required for maintenance of excitatory and inhibitory synapse numbers

Astrocytes control the formation and maintenance of synapses through secreted molecules^63–66^. However, whether astrocytic vesicular exocytosis is involved in astrocytes’ synaptogenic functions remains enigmatic. Because we found Vamp2 to be expressed in mouse cortical astrocytes **(Figure 5)**, we investigated its role in cortical astrocytes’ synaptogenic ability.

To do so, we deployed the GEARBOCS-GeneTrap method to KO astrocytic Vamp2 with concurrent mCherry-labeling of KO astrocytes. We retro-orbitally injected AAV-GEARBOCS-Vamp2-GeneTrap **(Figure 6A).** To determine if astrocytic Vamp2 is required to maintain proper synaptic connectivity, we quantified the numbers of excitatory and inhibitory synapses within the territories of Vamp2-GeneTrap and control astrocytes. For synapse quantification, we used an immunohistochemical method that takes advantage of the close proximity of pre- and post-synaptic proteins at synaptic junction^67,68^. Even though these proteins are at different cellular compartments (i.e., axons and dendrites, respectively), they appear to co-localize at synapses due to the resolution limit of light microscopy.

**Figure 6:**
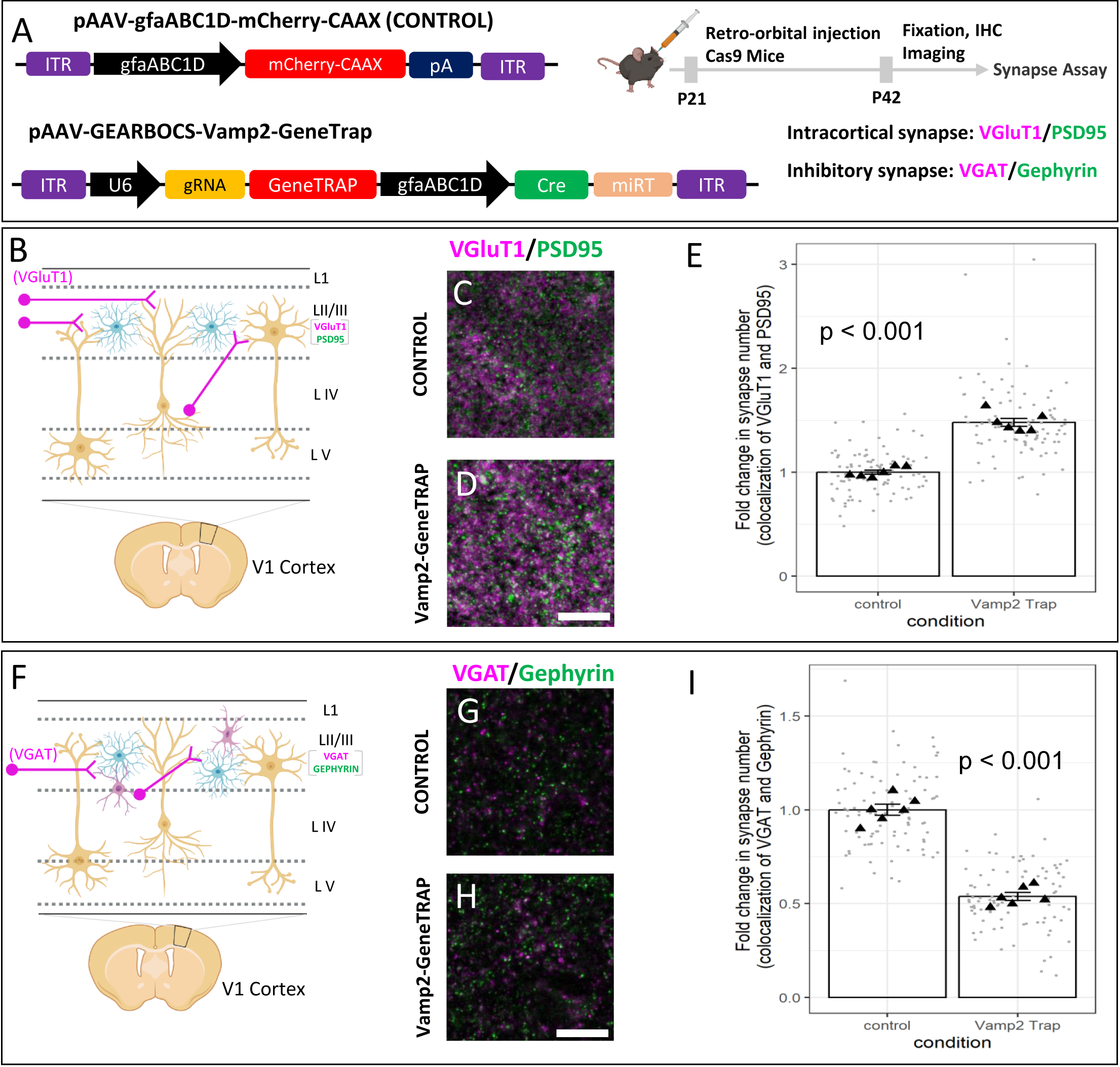
Astrocytic Vamp2 is required for regulating excitatory and inhibitory synapse numbers: **A).** Schematic of both AAV-GEARBOCS-*Vamp2*-GeneTrap and AAV-gfaABC1D-mCherry-CAAX plasmid and the experimental scheme of AAV injection and synapse number quantification assays. AAVs were generated with the AAV-PHP.eB capsid and retro-orbitally injected into the loxP-STOP-loxP Cas9 mice at P21 and the brains were collected at P42. For synapse assays we used presynaptic VGluT1 and postsynaptic PSD95 markers. **B).** Schematic showing layer II/III VGluT1 (magenta) intracortical presynaptic inputs in V1 cortex. **C-D)** Confocal immunofluorescence images of intracortical excitatory synapses marked as close apposition of VGluT1 (magenta) and PSD95 (green)**. E)** Quantification of synaptic density of intracortical co-localized puncta. 6 mice per condition. **F).** Schematic showing layer II/III VGAT (magenta) inhibitory presynaptic inputs in V1 cortex. **G-H)** Confocal immunofluorescence images of intracortical excitatory synapses marked as close apposition of VGAT (magenta) and Gephyrin (green)**. I)** Quantification of synaptic density of intracortical co-localized puncta. 6 mice per condition. p values were calculated using a linear mixed effects model to account for the multiple images taken from each animal. Error bars represent 1 standard error of the mean. Scale bars= 10μm.

To quantify excitatory cortical synapses, we labeled the brain sections with the excitatory intracortical presynaptic marker VGluT1^69^ and the postsynaptic marker PSD95. The inhibitory synapses were identified as the co-localization of presynaptic VGAT and postsynaptic Gephyrin. Interestingly, loss of Vamp2 in astrocytes caused a significant increase in the densities of intracortical (VGluT1/PSD95+) **(Figure 6B-E)** excitatory synapses when compared to control astrocytes. On the contrary, loss of Vamp2 caused a severe reduction in the density of VGAT/Gephyrin+ inhibitory synaptic puncta within the territories of Vamp2 KO astrocytes in layer 2/3 (**Figure 6F-L).**

Taken together, our results elucidate a previously unknown role for Vamp2 in mouse cortical astrocytes in maintaining the balance between the excitatory and inhibitory synapses. These findings suggest that Vamp2 controls the release of synapse-modifying factors. Our findings also demonstrate that GEARBOCS is a useful genome-editing tool for the investigation of astrocyte-mediated complex cellular and molecular mechanisms in CNS development and function.

## Discussion

To further our understanding of astrocyte biology in CNS homeostasis and function an efficient molecular tool for rapid *in vivo* genome-editing in mouse astrocytes is needed. In this study, we devised a HITI-based, single virus, CRISPR tool, GEARBOCS, to target mouse cortical astrocytes *in vivo* and demonstrated its multiple applications in genetic manipulation of astrocytes.

GEARBOCS has several advantages and applications. First, GEARBOCS is a single AAV vector capable of carrying all the essential components required for *in vivo* genome editing when used in conjunction with a Cre-dependent Cas9 transgenic mouse line. The single vector GEARBOCS strategy ensures the simultaneous and efficient delivery of all components to the astrocytes. Second, GEARBOCS allows for multiple genome-editing strategies including gene knock out or knocking in reporters or tags to determine astrocytic protein expression and localization for genes of interest. Furthermore, the GEARBOCS-mediated GeneTrap strategy can be used to sparsely label and visualize the KO astrocytes to study their morphology and synapse interactions. Importantly, GEARBOCS can accommodate donor sizes up to 2kb. The donors can be interchanged in GEARBOCS to generate KO, TagIn and GeneTrap options for a gene of interest quickly. Third, GEARBOCS, when used with AAV capsids such as PHP.eB, efficiently and broadly targets astrocytes across the CNS after non-invasive retro-orbital injections. The high efficacy of astrocytic transduction provides a quick cell-type specific gene manipulation capability. Noninvasive AAV delivery greatly reduces the risk of causing reactive astrocytosis, which is often observed following direct injection of viruses into the brain^70^.

In this study, we used GEARBOCS technique to determine the necessity of astrocytic Sparcl1 for thalamocortical synapse formation and maintenance during and after the critical period for ocular dominance plasticity. Previous studies have shown that Sparcl1 is required for proper synapse formation and that overexpression of Sparcl1 specifically in astrocytes is sufficient to rescue this phenotype. However, determining the necessity of astrocytic Sparcl1 required a tool like GEARBOCS to specifically KO Sparcl1 only in astrocytes, since other cell types could serve as a compensatory source of Sparcl1. Our experiments found that both injection of GEARBOCS-Sparcl1-KO at P21 followed by P42 collection and injection at P42 followed by P62 collection resulted in a reduction VGluT2/PSD95 thalamocortical synapse numbers, without altering VGlut1/PSD95 intracortical synapse numbers. These results show that astrocytes serve as the primary source of Sparcl1 in the mouse visual cortex to maintain thalamocortical synapses.

We also investigated the expression of Vamp2 in astrocytes *in vivo* using GEARBOCS. Calcium-regulated vesicular exocytosis is a key feature of neuronal synapses and is mediated by the molecular machinery of the SNARE complex including Vamp2^36,37^. Several studies suggested that Vamp2, and other SNARE complex proteins, are involved in secretion of small neuroactive molecules and proteins from astrocytes^59,71,72^. However, it has been difficult and controversial to provide evidence for Vamp2 expression and function in astrocytes^38,39,61^. In this study, we used GEARBOCS to delineate the expression of Vamp2 in mouse cortical astrocytes *in vivo*. We were able to detect reporter expression under the control of Vamp2 endogenous promoter via GeneTrap method and we found vesicle-like HA-tag staining within Vamp2-TagIn astrocytes. These data provide further evidence that Vamp2 is expressed by astrocytes *in vivo*.

GEARBOCS-mediated endogenous tagging and visualization of proteins such as Vamp2 circumvents the challenges and limitations of antibody-based immunolabeling of proteins in astrocytes *in vivo*. Thus, GEARBOCS facilitates the study of essentially any gene of interest in astrocytes, even in the absence of specific antibodies or conditional alleles. Moreover, this CRISPR tool allows for the determination of astrocyte-specific localization and distribution of proteins, even when the protein of interest, such as Vamp2, is more abundant in another CNS cell type, such as neurons. In addition to these advantages, GEARBOCS can be used to replace the strategy of ectopic expression of tagged proteins which may alter their localization and function in astrocytes.

Astrocytes form an integral part of the synapse and control excitatory and inhibitory synaptogenesis through the secretion of synapse-modifying proteins^17,20,73^. Moreover, astrocytes have been proposed to participate in the regulation of neural circuits by secreting neuroactive small molecules such as D-serine and ATP that modify synaptic activity^72,74^. Because Vamp2 is well established to control synaptic neurotransmitter release in neurons^36^, as well as protein secretion in non-neuronal cell types^75^, we postulated that loss of Vamp2 will alter the regulation of excitatory and inhibitory synapse numbers locally. In agreement with this possibility, we found that loss of Vamp2 in astrocytes after the end of synaptogenic period (P21) significantly altered synapse numbers. Loss of astrocytic Vamp2 increased intracortical excitatory synapse numbers in the visual cortex, whereas the number of inhibitory synapses were reduced. These observations indicate that Vamp2-mediated secretion is required for maintaining the excitation/inhibition balance.

Astrocytes undergo local and global Ca^2+^-transients, albeit at much slower timescales than neurons. The astrocytic Ca^2+^-transients occur in response to patterned neuronal activity and neuromodulators^76,77^. Manipulation of astrocytic calcium through opto- and chemo-genetic tools result in strong changes in synapse activity and animal behavior^78^. Vamp2 controls Ca^2+^-dependent exocytosis in neurons^79,80^. Our results suggest that Vamp2 could also mediate astrocytic exocytosis of synapse-modifying molecules. Thus, our finding reveals a rich and unexplored aspect of astrocytic-secretion and provides the rationale for future studies to study Vamp2 function in astrocytes *in vivo*.

In summary, our results show that GEARBOCS is an effective and multifunctional gene-editing tool that offers a quick and efficient way to investigate astrocyte biology at the cellular and molecular levels in mice. The simple design of GEARBOCS makes it a powerful and versatile ‘all-in-one’ CRISPR tool for both basic and translational research. Thus, future studies using GEARBOCS will likely provide new insights into the roles of astrocytes in the pathophysiology of neurodevelopmental and neurodegenerative diseases.

## Limitations

GEARBOCS TagIn strategy enables both N and C-terminal tagging. However, an important caveat with introducing N-terminal tags is; it could lead to the gRNA-mediated KO of the gene if the donor integration does not happen. Similarly, GeneTrap strategy targets donors with reporters close to the START codon and in the absence of donor integration the Cas9+ astrocytes could be KOs. Therefore, the neighboring unlabeled astrocytes, which express Cas9, should not be considered WT.

In the future, the GEARBOCS method can be further developed and improved in several ways. GEARBOCS is a modular tool; thus, we can swap the promoters of gRNA and the Cre recombinase in a species- or cell type-specific manner to achieve broader applicability. For example, GEARBOCS can be modified to utilize smaller Cas9 homologs such as SaCas9^81^ under the gfaABC1D promoter instead of driving Cas9 expression via Cre recombinase. This would eliminate the need for using Cas9 transgenic mouse lines and permit the use of GEARBOCS in other species.

The efficiency of GEARBOCS-mediated KO, TagIn and GeneTrap methods depend on many factors. The most important one is the effectiveness and specificity of the gRNA, which is heavily influenced by the accessibility of the targeted genomic locus. Off-target activity of gRNAs and imprecise insertion of donors pose significant challenges in CRISPR-mediated genome editing^82^. Therefore, careful validation of the gRNAs for the genes of interest are critical for experimental success. In this study, we used guides which were previously validated or validated the guides for Sparcl1 using othologous methods such as by using 3T3-Cas9 cells^12,62^. It is critical that future studies using GEARBOCS with newly developed guides should screen for off-target activity or precise insertion of tags at endogenous loci^83^.

## Methods

### Resource Availability

#### Lead contact

Further information and requests for resources and reagents should be directed to the lead contact, Cagla Eroglu.

## Materials availability

Plasmids for all viral constructs produced in this study will be deposited on Addgene and made available upon publication.

## Data and code availability

- Raw data and analysis can be found can be accessed on Zenodo at the following DOI: 10.5281/zenodo.13887806.
- R code related to synapse quantification in this paper is available at https://github.com/Eroglu-Lab/gearbocs_analysis and on Zenodo at the following DOI: 10.5281/zenodo.13312831.
- Any additional information can be requested from the lead contact.

## Experimental model and subject details

### Animals

All mice experiments were carried out under a protocol approved by the Duke University Institutional Animal Care and Use Committee (IACUC) in accordance with US National Institutes of Health guidelines. All mice were housed in the Duke Division of Laboratory Animal Resources (DLAR) facility and follow typical day/night conditions of 12-hours cycles. Both B6J.129(B6N)-Gt (ROSA)26Sortm1(CAG-cas9*,EGFP)Fezh/J (JAX stock #026175) and B6.129-Igs2tm1(CAG-cas9*)Mmw/J (JAX stock #027632) mice were purchased from Jackson laboratory. AAV retro-orbital injections were performed at either P21 or P42, and brains collected at P42 or P65 respectively. Sex-matched littermate pairs (both male and female) were randomly assigned to experimental groups for all experiments.

## Method Details

### Plasmids and CRISPR guides

To generate GEARBOCS, Cre expression cassette was cloned first into the pZac2.1-GfaABC1D-Lck-GCaMP6f (A gift from Dr. Baljit Khakh; Addgene plasmid #52924) by replacing Lck-GCaMP6f. U6 expression cassette along with the donor insertion sites (DIS) were synthesized as gBlocks (IDT) and cloned upstream of gfaABC1D promoter. The 4x6T microRNA targeting cassette from AAV-GfaABC1D-Cre-4x6T (Addgene plasmid #196410) was synthesized as a gBlock (IDT) and cloned between the Cre sequence and SV40 polyA signal. pAAV-gfaABC1D-mCherry-CAAX was generated by cloning mCherry-CAAX into pZac2.1-GfaABC1D-Lck-GCaMP6f by replacing Lck-GCaMP6f. All the gRNAs used in this study were cloned into the Sap1 site of GEARBOCS and the donors were cloned between the Sal1 and BamH1 site in the DIS. To make the GEARBOCS donors, mCherry donors were PCR amplified and HA donors were oligo annealed with overhangs to clone into the DIS. All generated plasmids were confirmed by Sanger sequencing protocol. The sgRNA sequences used in this study were as follows:

Sparcl1: KO and gene trap 5’-GTGCGCCTTGGGAACCGCTGTGG-3’,

TagIN C-term 5’-CGAGTTATGCAGTGTTCCATGGG-3’,

TagIN N-term 5’-CGCTACAGTCCCCATTGCCGGGG-3’.

VAMP2: Gene trap and TagIN N-term 5’-GGCCGGGGCGGCAGGCGGGACGG-3’

For a detailed protocol for designing guide RNAs and cloning into the GEARBOCS vector see https://www.protocols.io/view/gearbocs-protocols-36wgqnq15gk5/v3.

## AAV production and purification

Purified AAVs were produced as previously described (Uezu et al., 2016). Briefly, HEK293T cells grown on 150mm dishes were transfected with GEARBOCS plasmid, helper plasmid pAd-DeltaF6 (Addgene Plasmid #112867) and the serotype plasmid AAV PHP.eB (Addgene Plasmid #103005). After three days, cell lysates were prepared with 15mM NaCl, 5mM Tris-HCl, pH 8.5 followed by three repeats of freeze-thaw cycles. The cell lysates were centrifuged for 30min at 4000rpm to collect the supernatant. Benzonase-treated supernatant (50U/ml, 30 min at 37°C) was added to an Optiprep density gradient (15%, 25%, 40% and 60%) for ultracentrifugation at 60,000 rpm for 1.5hr using a Beckman Ti-70 rotor. AAV fraction was collected from the gradient and concentrated along with multiple washes with DPBS in a 100 kDa filtration unit. AAV titers were quantified by qPCR based on SYBR green technology using primer pair targeting AAV2 ITR (Aurnhammer et al., 2012). For a detailed protocol see https://www.protocols.io/view/gearbocs-protocols-36wgqnq15gk5/v3.

## AAV injection and tissue preparation

P21 or P42 Cas9 or Cas9-EGFP mice placed in a stereotaxic frame were anesthetized through inhalation of 1.5% isoflurane gas. 10μl of purified AAVs (titer of ∼1 x 10^12^ GC/ml) was intravenously injected into the retro-orbital sinus. After 3 weeks at P42 or P65, mice were anesthetized with 200 mg/kg Tribromoethanol (Avertin) and transcardially perfused with ice-cold TBS/Heparin and 4% paraformaldehyde (PFA) at room temperature (RT). Harvested brains were post-fixed overnight in 4% PFA, cryoprotected in 30% sucrose and the brain blocks were prepared with O.C.T. (TissueTek) to store at −80°C. 25µm thick brain sections were obtained through cryo-sectioning using a Leica CM3050S (Leica, Germany) vibratome and stored in a mixture of TBS and glycerol at −20°C for further free-float antibody staining procedures. For a detailed protocol see https://www.protocols.io/view/gearbocs-protocols-36wgqnq15gk5/v3.

## HEK293T cell culture

For AAV production, HEK293T (ATCC CRL-11268) cells were cultured in DMEM high glucose medium (Thermo Fisher #11960044) supplemented with 2mM L-Glutamine, 10% fetal bovine serum, 100U/ml Penicillin-Streptomycin, and 1 mM sodium pyruvate. Cells were incubated at 37°C in 5% CO2 and passaged by trypsin/EDTA digestion upon reaching ∼95% confluency. Cells were transfected with PEI Max (Polysciences) when reaching 60-80% confluency. For a detailed protocol see https://www.protocols.io/view/gearbocs-protocols-36wgqnq15gk5/v3.

## In-Vitro CRISPR Guide Validation

Initial guide RNA screening was done utilizing NIH3T3/Cas9 (Calibre Scientific #CBIO-AKR-5104) cells. The cells were cultured in DMEM high glucose medium (Thermo Fisher #11960044) supplemented with 2mM L-Glutamine, 10% fetal bovine serum, 100U/ml Penicillin-Streptomycin, and 1 mM sodium pyruvate. Cells were incubated at 37°C in 5% CO2, and passaged by trypsin/EDTA digestion upon reaching ∼90% confluency. For guide screening, 250k NIH3T3/Cas9 cells were seeded into each well of a 6-well plate. 24 hours after seeding, the cells were transfected with the GEARBOCS plasmid containing the guide RNA to be validated. 48 hours after transfection, the cells were collected via scraping and the genomic DNA was isolated using a Qiagen DNeasy kit (Qiagen #69504). The region targeted by the guide was then amplified via PCR and purified via gel extraction. The PCR product was then analyzed via Sanger sequencing. Successful guides were determined by the detection of the insert DNA for TagIN or gene-trap constructs. For knockout constructs, success was determined by a drop in read quality after the targeted cut site due to a loss of consensus after the successful introduction of indel mutations. For a detailed protocol see https://www.protocols.io/view/gearbocs-protocols-36wgqnq15gk5/v3.

## Immunohistochemistry

For immunohistochemistry, frozen tissue sections were washed three times in 0.2% TBST (0.2% Triton X-100 in 1x TBS). The sections were blocked and permeabilized with a blocking buffer (5% normal goat serum in 0.2% TBST) for 1hr at RT followed by an overnight incubation with the primary antibodies at 4°C. Primary antibodies were diluted in blocking buffer. After primary incubation, sections were washed in 0.2 % TBST and incubated with fluorescent secondary antibodies diluted in blocking buffer for 2-3 hours at RT. Tissue sections were washed in TBST and mounted on glass slides with Vectashield with DAPI (Vector Laboratories, CA). Primary antibodies used for immunohistochemistry were listed as following: Rat anti-HA (Roche/Sigma #11867423001), Chicken anti-GFP (Aves Labs #GFP-1020), Rabbit anti-RFP (Rockland #600-401-379), Rabbit anti-Sox9 (Millipore #AB5535), Rabbit anti-Vamp2 (Proteintech # 10135-1-AP), Mouse anti-NeuN (Millipore # MAB377), Mouse anti-Olig2 (Millipore #MABN50), Chicken anti-mCherry (Aves Labs #MCHERRY-0020), Mouse anti-Vamp2 (Synaptic Systems # 104211), Guinea pig anti-VGAT (Synaptic Systems #131004), Rabbit anti-PSD95 (Thermo Fisher Scientific #51–6900), Guinea pig anti-VGLUT1 (Millipore # AB5905), Guinea pig anti-VGLUT2 (Synaptic Systems #135 404), Mouse anti-gephyrin (Synaptic Systems #147–011), and Rat anti-hevin (12-155)^84^. Goat Alexa Fluor Secondary antibodies (Invitrogen) used for immunohistochemistry were listed as following: Rabbit 488 (#A-11034), Guinea pig 647(#A-21450), Chicken 488(#A-11039), Chicken 594 (#A-11042), Rabbit 594 (#A-11037), Rat 594(#A-11007), Mouse 594(#A-21125), Mouse 568 (#A-21134), Mouse 488 (#A-21121), Rat 568 (#A-11077) and Mouse 647 (#A-21240). To avoid excessive background staining, isotype specific secondary antibodies were used for primary antibodies produced in mice. For a more detailed protocol see https://www.protocols.io/view/synapse-staining-ihc-vglut1-and-psd95-mouse-brain-6qpvr3x4zvmk/v1.

## Synaptic Puncta Colocalization Analysis

Confocal images of pre- and postsynaptic puncta were prepared for analysis using ImageJ (https://imagej.nih.gov/ij/) to convert the raw images into RGB type images with the presynaptic marker (VGlut1/VGlut2 or VGAT) in the red channel, the postsynaptic marker in the green channel (PSD95 or Gephyrin), and the astrocyte marker (mCherry) in the blue channel. Images were then Z-projected to make one projection for every 3 images, representing 1 um of depth in our imaging setup. Colocalizations between the red and green channels of these Z-stack images were then quantified using the Puncta Analyzer FIJI plugin as described in Ippolito and Eroglu 2010^68^. A linear mixed effects model was used to test the effect of experimental condition on the synapse counts while accounting for the multiple images from each animal and the paired imaging design (staining and imaging were done for pairs of control and experimental animals at the same time). More details for these statistical analyses can be found in our R scripts available at https://github.com/Eroglu-Lab/gearbocs_analysis.

## Statistical Analysis

For data collection, brains from the healthy mice in each experimental group were collected and processed. All data are represented as mean ± standard error of the mean and P < 0.05 were considered to indicate statistical significance. Exact P-values are noted in the figure/figure legends for each experiment. Statistical analyses (other than the Synaptic Puncta Colocalization Analysis above) were performed with JMP Genomics Pro 17.0 software, and the graphs were generated with GraphPad Prism version 9 (GraphPad Software). Statistical analyses of the quantified data were performed using either Nested analysis of variance (ANOVA) or Nested t-test. Sample size for each experiment and statistical method of analysis are indicated in the figure legend for each experiment.

## Supporting information

Resources Table

## Acknowledgments

We thank Drs. Dolores Irala, Francesco P. Ulloa Severino, Shiyi Wang, Kristina Sakers and other Eroglu lab members for critical feedback about the manuscript. We thank Adam Okinaga for excellent technical assistance. This work was supported by Chan Zuckerberg Initiative, Neurodegeneration Challenge Network Collaborative grant (2018-191999) and two NIH BRAIN Initiative Grants (R01-DA047258 and U19-NS123719) to CE and by the joint efforts of MJFF and the Aligning Science Across Parkinson’s (ASAP) initiative. MJFF administers the grant [ASAP-020607 to CE] on behalf of ASAP and itself Michael J Fox Foundation. JTS and NB are supported by NIH funding F31NS134252 and 5T32HD040372-19 respectively. Cagla Eroglu is an HHMI Investigator. This article is subject to HHMI’s Open Access to Publications policy. HHMI lab heads have previously granted a nonexclusive CC BY 4.0 license to the public and a sublicensable license to HHMI in their research articles. Pursuant to those licenses, the author-accepted manuscript of this article can be made freely available under a CC BY 4.0 license immediately upon publication.

## Author Contributions

Conceptualization, DSB and CE; Methodology, DSB, CXT, JTS; Investigation, DSB, NB, KD, CXT, JTS; Formal analysis, DSB, NB, CXT and JTS; Software, JTS; Writing – original draft, DSB and CE; Writing – Review & Editing, DSB, JTS, NB, CXT, LB, and CE, Funding Acquisition, CE.

