## Supplemental Information for "GEARBOCS: An Adeno Associated Virus Tool for *In Vivo* Gene Editing in Astrocytes"

**Supplemental Figure 1** **Sivadasan Bindu *et al.***

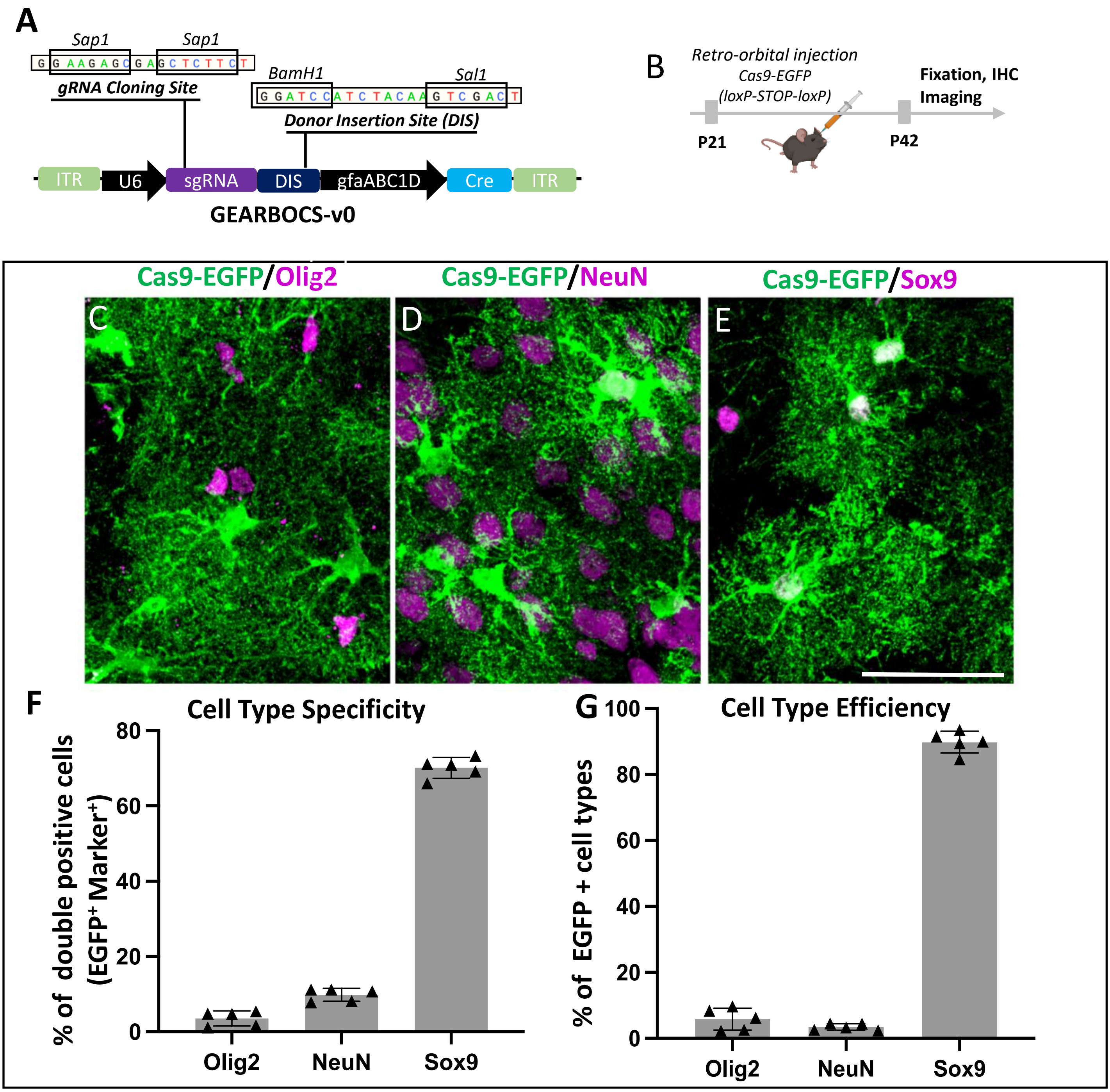

**Figure S1: GEARBOCS-v0 expresses Cre in astrocytes, with some off-target activity in neurons**

**A)** Strategy for GEARBOCS-v0 including a CRISPR gRNA cloning site and donor insertion site (DIS) in the same AAV viral construct. **B)** Timeline of GEARBOCS injections. Includes P21 retroorbital AAV injection followed by P42 animal collection for immunohistochemistry. **C-E)** Cas9-EGFP signal in GEARBOCS-v0 injected mouse visual cortex with Olig2, NeuN, or Sox9 staining to assess cell type-specific expression in oligodendrocytes, astrocytes, or neurons respectively. Scale bar represents 20μm. **F)** Plot showing the cell type distribution out of all of the EGFP positive cells (Specificity) **G)** Quantification of what percentage of cells in a given cell type were EGFP+ (Efficiency). (F&G: n=5 animals, Error bars represent one standard error of the mean).

### Supplemental Figure 2

Sivadasan Bindu *et al.*

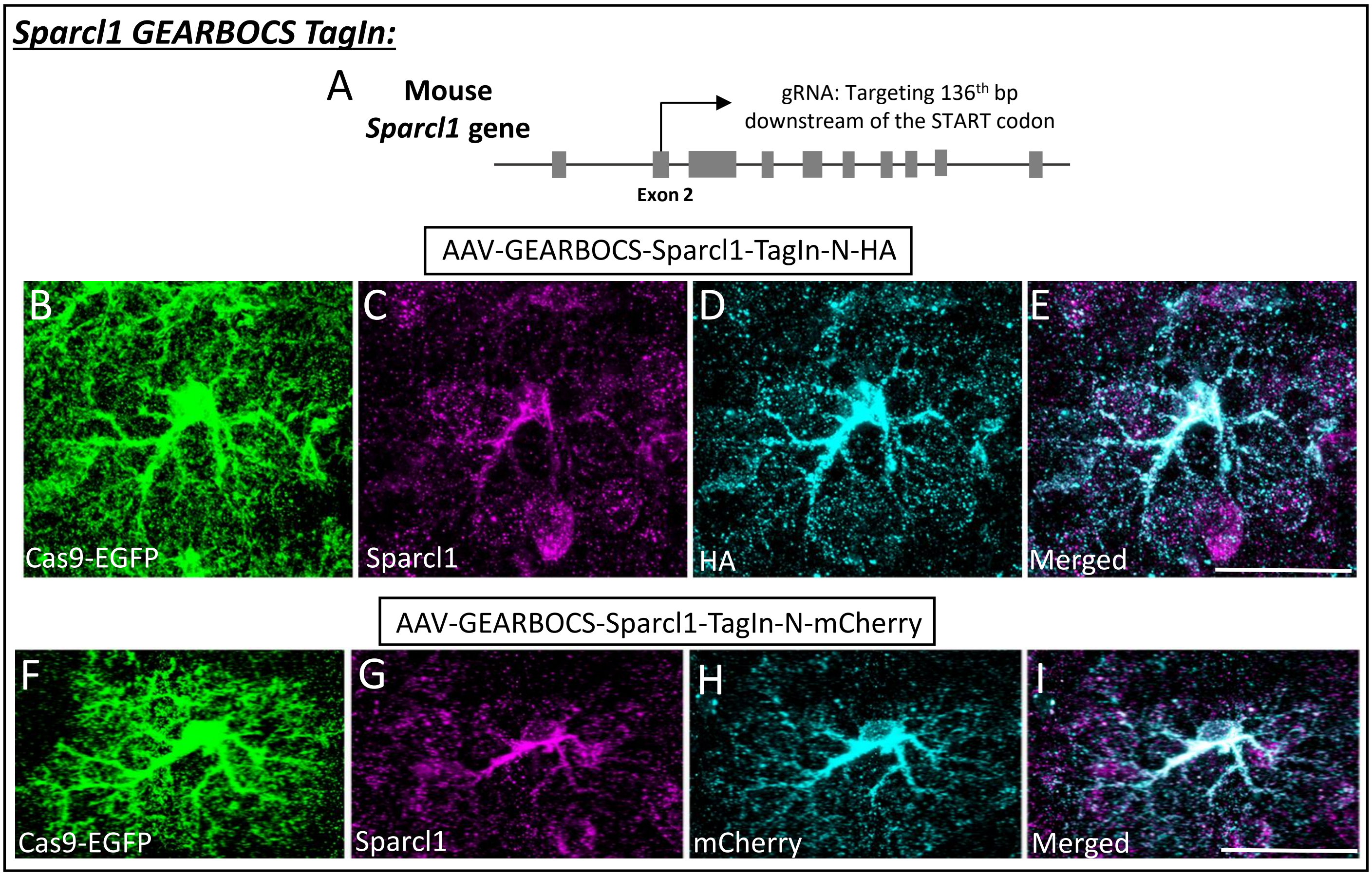


J

CRISPR Validation-Genomic Integration

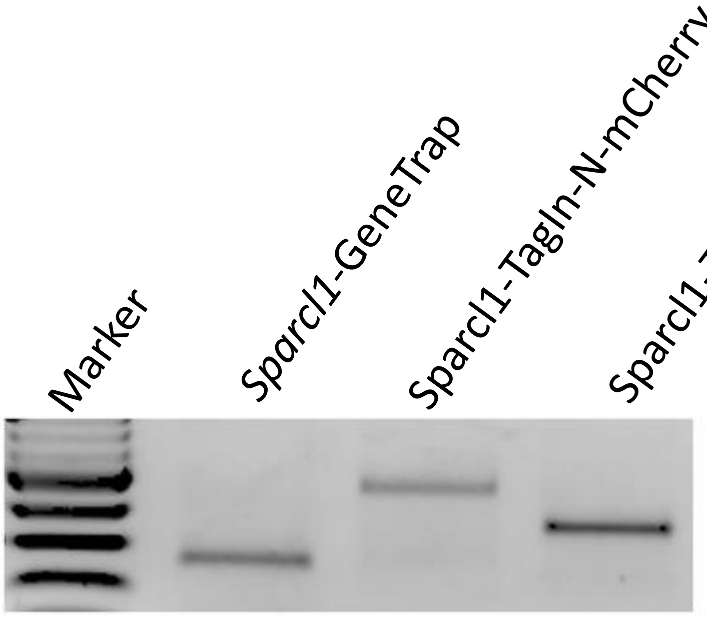

**Figure S2: GEARBOCS can label astrocytic Sparcl1 at the N-terminus**

**A)** N-terminus targeting CRISPR gRNA for Sparcl1 used to AAV-GEARBOCS-Sparcl1-TagIn-N-HA and AAV-GEARBOCS-Sparcl1-TagIn-N-mCherry. **B-E)** Cas9-EGFP, Sparcl1, and N-terminal HA tag immunohistochemistry in a mouse visual cortex astrocyte. **F-I)** Cas9-EGFP, Sparcl1, and N-terminal mCherry tag immunohistochemistry in a mouse visual cortex astrocyte. **J)** Validation of genomic integration of GeneTrap, TagIn-N, and TagIn-C strategies targeting Sparcl1 through PCR amplification of genomic DNA from the edited region in edited 3T3 cells. Scale bars=20um.

### Supplemental Figure 3

Sivadasan Bindu *et al.*

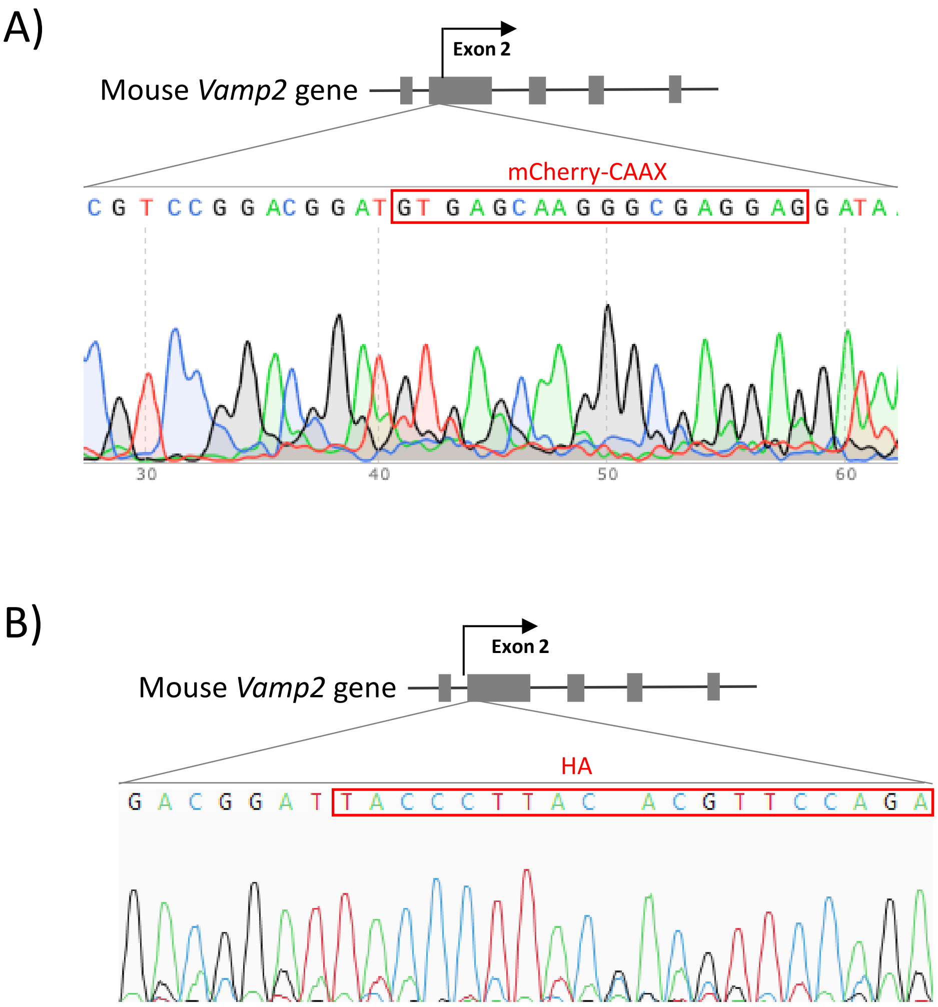

**Figure S3: GEARBOCS can label astrocytic Vamp2 at the N-terminus**

**A)** Sanger sequencing validation of Vamp2 gene editing for mCherry-CAAX TagIn at the N-terminus using Sanger sequencing of genomic DNA from edited 3T3 cells. **B)** Sanger sequencing validation of Vamp2 gene editing for HA TagIn at the N-terminus using Sanger sequencing of genomic DNA from edited 3T3 cells.
