## Supplementary material for "GEARBOCS: An Adeno Associated Virus Tool for *In Vivo* Gene Editing in Astrocytes": Resources Table

| **REAGENT or RESOURCE** | SOURCE | IDENTIFIER |
|---|---|---|
| **Antibodies** | | |
| Rat-anti-HA | Roche | Cat# 11867423001, RRID:AB_390918 |
| Chicken-anti-GFP | Aves Labs | Cat# GFP-1020, RRID:AB_10000240 |
| Rabbit-anti-RFP | Rockland | Cat# 600-401-379, RRID:AB_2209751 |
| Rabbit-anti-Sox9 | Millipore | Cat# AB5535, RRID:AB_2239761 |
| Rabbit-anti-Vamp2 | Proteintech | Cat# 10135-1-AP, RRID:AB_2256918 |
| Mouse-anti-NeuN | Millipore | Cat# MAB377, RRID:AB_2298772 |
| Mouse-anti-Olig2 | Millipore | Cat# MABN50, RRID:AB_10807410 |
| Chicken-anti-mCherry | Aves Labs | Cat #MCHERRY-0020, RRID:AB_2910557 |
| Mouse-anti-Vamp2 | Synaptic Systems | Cat# 104 403, RRID:AB_2864782 |
| Mouse-anti-VGAT | Synaptic Systems | Cat# 131 004, RRID:AB_887873 |
| Rabbit-anti-PSD95 | Thermo Fisher Scientific | Cat# 51-6900, RRID:AB_2533914 |
| Guinea Pig-anti-VGLUT1 | Millipore | Cat# AB5905, RRID:AB_2301751 |
| Guinea Pig-anti-VGLUT2 | Synaptic Systems | Cat# 135 404, RRID:AB_887884 |
| Mouse-anti-Gephyrin | Synaptic Systems | Cat# 147 011, RRID:AB_887717 |
| Rabbit Alexa Fluor™ 488 | Invitrogen | Cat# A-11034 RRID: AB_2576217 |
| Guinea pig Alexa Fluor™ 647 | Invitrogen | Cat# A-21450 RRID: AB_2535867 |
| Chicken Alexa Fluor™ 488 | Invitrogen | Cat# A-11039 RRID: AB_2534096 |
| Chicken Alexa Fluor™ 594 | Invitrogen | Cat# A-11042 RRID: AB_2534099 |
| Rabbit Alexa Fluor™ 594 | Invitrogen | Cat# A-11037 RRID: AB_2534095 |
| Rat Alexa Fluor™ 594 | Invitrogen | Cat# A-11007 RRID: AB_10561522 |
| Mouse Alexa Fluor™ 594 | Invitrogen | Cat# A-21125 RRID: AB_2535767 |

| | | |
|---|---|---|
| Mouse Alexa Fluor™ 568 | Invitrogen | Cat# A-21134 RRID: AB_2535773 |
| Mouse Alexa Fluor™ 488 | Invitrogen | Cat# A-21121 RRID: AB_2535764 |
| Rat Alexa Fluor™ 568 | Invitrogen | Cat# A-11077 RRID: AB_2534121 |
| Mouse Alexa Fluor™ 647 | Invitrogen | Cat# A-21240 RRID: AB_2535809 |
| **Bacterial and virus strains** | | |
| One Shot™ Stbl3™ Chemically Competent E. coli | Invitrogen | Cat# C737303 |
| AAV-GEARBOCS-v0 | This Study | NA |
| AAV-GEARBOCS (with 4xmiRT) | This Study | NA |
| AAV-GEARBOCS-Sparcl1-KO | This Study | NA |
| AAV-GEARBOCS-Sparcl1-TagIn-C-mCherry | This Study | NA |
| AAV-GEARBOCS-Sparcl1-TagIn-N-mCherry | This Study | NA |
| AAV-GEARBOCS-Sparcl1-TagIn-N-HA | This Study | NA |
| AAV-GEARBOCS-Sparcl1_GeneTRAP | This Study | NA |
| AAV-GEARBOCS-*Vamp2*-GeneTRAP | This Study | NA |
| AAV-GEARBOCS-v0-*Vamp2*-TagIN-HA | This Study | NA |
| AAV-GfaABC1D-mCherry-CAAX | This Study | NA |
| **Chemicals, Peptides, and Recombinant Proteins** | | |
| PEI MAX® | Polysciences | Cat#24765 |
| Optiprep | Sigma | Cat# D1556 |
| Pen/Strep | GIBCO | Cat# 15140 |
| Sodium Pyruvate | GIBCO | Cat# 11360-070 |
| L-Glutamine | GIBCO | Cat# 25030-081 |
| DMEM | GIBCO | Cat# 11960044 |

| | | |
|---|---|---|
| DPBS | GIBCO | Cat# 14190144 |
| Benzonase | Novagen | Cat#70664 |
| Goat Serum | GIBCO | Cat#16210064 |
| Triton™ X-100 Surfact-Amps™ Detergent Solution | Thermo Scientific | Cat#28314 |
| Fetal Bovine Serum | Sigma | Cat#F4135 |
| 2,2,2 tribromoethanol (Avertin) | Sigma | Cat# T48402-25G |
| 2-methyl-2-butanol | Sigma | Cat# 152463-250mL |
| Opti-MEM | GIBCO | Cat# 31985070 |
| X-tremeGENE HP DNA Transfection Reagent | Sigma-Aldrich | Cat# 6366244001 |
| PFA 16% | Electron Microscopy Sciences | Cat# 15710 |
| Tissue-Tek O.C.T. Compound | Sakura Finetek | Cat# 4583 |
| Heparin | Sigma | Cat# H3149-10KU |
| **Critical Commercial Assays** | | |
| Endo-Free Maxi Prep Kit | QIAGEN | Cat# 12362 |
| QIAprep Spin Miniprep Kit | QIAGEN | Cat# 27106 |
| QIAquick Gel Extraction Kit | QIAGEN | Cat# 28704 |
| Vivaspin™ ultrafiltration spin columns | Cytiva | Cat# 28932363 |
| Zero Blunt™ TOPO™ PCR Cloning Kit | Invitrogen | Cat# 450031 |
| In-Fusion® Snap Assembly Master Mix | Takara | Cat# 638948 |
| Fast SYBR™ Green Master Mix | Applied Biosystems™ | Cat#4385612 |
| Phusion® High-Fidelity PCR Kit | NEB | Cat# E0553L |
| DNeasy Blood & Tissue Kit | QIAGEN | Cat# 69504 |

| | | |
|---|---|---|
| **Experimental models: cell lines** | | |
| HEK293T | ATCC | CRL-11268, RRID: CVCL_1926 |
| NIH3T3/Cas9 Cell Line | Calibre Scientific | CBIO-AKR-5104, RRID: |
| **Experimental models: organisms/strains** | | |
| B6J.129(B6N)-Gt (ROSA)26Sortm1(CAG-cas9*,-EGFP)Fezh/J | Jackson Laboratory | Cat#026175 RRID: IMSR_JAX: 026175 |
| B6.129-Igs2tm1(CAG-cas9*)Mmw/J | Jackson Laboratory | Cat#027632, RRID: IMSR_JAX: 027632 |
| **Oligonucleotides** | | |
| *Sparcl1* gRNA | This Study | See Methods |
| *Vamp2* gRNA | This Study | See Methods |
| **Equipment** | | |
| Beckman Ti70 rotor | | |
| Beckman preparative ultracentrifuge | | |
| U-100 Insulin syringes | BD | Cat# 324702 |
| **Software and algorithms** | | |
| GraphPad Prism 9.4.1 | GraphPAD | https://www.graphpad.com/scientific-software/prism/ RRID: SCR_002798 |
| ImageJ | NIH | https://imagej.nih.gov/ij/ RRID: SCR_003070 |
| Puncta Analyzer (version 2.0) | Ippolito and Eroglu, 2010 | https://github.com/toddstavish/puncta-analyzer RRID: SCR_025425 |
| Imaris 9.9.0 | BitPlane | https://imaris.oxinst.com/packages RRID: SCR_007370 |
| JMP® Pro 17 | SAS | https://www.jmp.com RRID: SCR_022199 |

| | | |
|---|---|---|
| R | The R Foundation | https://www.r-project.org/ RRID: SCR_001905 |
| CRISPick | Broad Institute | https://portals.broadinstitute.org/gppx/crispick/public |
| **Recombinant DNA** | | |
| AAV-PHP.eB_capsid | Addgene | RRID:Addgene_103005 |
| pAD-ΔF6 | Addgene | RRID:Addgene_112867 |
| pZac2.1-GfaABC1D-Lck-GCaMP6f | Addgene | RRID: Addgene_52924 |
| pGEARBOCS-v0 | This Study | RRID:Addgene_196495 |
| pGEARBOCS (with 4xmiRT) | This Study | RRID:Addgene_218181 |
| pGEARBOCS-Sparcl1-KO | This Study | RRID:Addgene_218182 |
| pGEARBOCS-Sparcl1-TagIn-C-mCherry | This Study | RRID:Addgene_218183 |
| pGEARBOCS-Sparcl1-TagIn-N-mCherry | This Study | RRID:Addgene_218186 |
| pGEARBOCS-Sparcl1-TagIn-N-HA | This Study | RRID:Addgene_218185 |
| pGEARBOCS-Sparcl1_GeneTRAP | This Study | RRID:Addgene_218184 |
| pGEARBOCS-*Vamp2*-GeneTRAP | This Study | RRID:Addgene_218187 |
| pGEARBOCS-v0-*Vamp2*-TagIN-HA | This Study | RRID:Addgene_196494 |
| pGfaABC1D-mCherry-CAAX | This Study | RRID:Addgene_218189 |
| **Other** | | |
| Centrifugation tubes for freeze-thawing: with centristar cap | Corning | Cat# 430828 |
| Ultracentrifuge sealing tubes, optiseal | Beckman | Cat# 361625 |
